# Recruitment of CTCF to the SIRT1 promoter after Oxidative Stress mediates Cardioprotective Transcription

**DOI:** 10.1101/2024.05.17.594600

**Authors:** Tobias Wagner, Priyanka Priyanka, Rudi Micheletti, Meyer J. Friedman, Sreejith J. Nair, Amir Gamliel, Havilah Taylor, Xiaoyuan Song, Miook Cho, Soohwan Oh, Wenbo Li, Jeehae Han, Kenneth A. Ohgi, Madeline Abrass, Agnieszka D’Antonio-Chronowska, Matteo D’Antonio, Helen Hazuda, Ravindranath Duggirala, John Blangero, Sheng Ding, Carlos Guzmann, Kelly A. Frazer, Aneel K. Aggarwal, Alice E. Zemljic-Harpf, Michael G. Rosenfeld, Yousin Suh

**Affiliations:** School and Department of Medicine, University of California San Diego, La Jolla, CA 92093, USA; Cellular and Molecular Medicine, Department of Medicine, University of California San Diego, La Jolla, CA 92093, USA; Haya Therapeutics, 3210 Merryfield Row, San Diego, CA 92121, USA; Department of Oncology, Georgetown University, Washington, DC 20057, USA; School of Life Science, University of Science & Technology of China, Hefei Anhui 230027, P.R. China; Department of Genetics, Albert Einstein College of Medicine, Bronx, NY 10461, USA; Korea University Sejong Campus, Department of Pharmacy, Sejong 30019, South Korea; Department of Biochemistry and Molecular Biology, McGovern Medical School, University of Texas Health Science Center, Houston, TX 77030, USA; Department of Obstetrics and Gynecology, Columbia University, New York, NY 10461, USA; Institute of Genomic Medicine and Department of Pediatrics, University of California San Diego, La Jolla, CA 92093, USA; UTRGV School of Medicine, Brownsville, TX 78520, USA; Department of Health and Behavioral Sciences, Texas A&M University - San Antonio, TX 78224, USA; Gladstone Institute of Cardiovascular Disease, Gladstone Institutes, San Francisco, CA 94158, USA; Department of Bioengineering, Graduate Program in Bioinformatics & Systems Biology; UC San Diego, La Jolla, CA 92093, USA; Department of Pharmacological Sciences, Icahn School of Medicine at Mount Sinai, New York, NY 10029, USA; Department of Anesthesiology, University of California, San Diego, La Jolla, CA 92093, USA; Veterans Affairs San Diego Healthcare System, San Diego, CA 92093, USA

## Abstract

Because most DNA-binding transcription factors (dbTFs), including the architectural regulator CTCF, bind RNA and exhibit di-/multimerization, a central conundrum is whether these distinct properties are regulated post-transcriptionally to modulate transcriptional programs. Here, investigating stress-dependent activation of *SIRT1,* encoding an evolutionarily-conserved protein deacetylase, we show that induced phosphorylation of CTCF acts as a rheostat to permit CTCF occupancy of low-affinity promoter DNA sites to precisely the levels necessary. This CTCF recruitment to the SIRT1 promoter is eliciting a cardioprotective cardiomyocyte transcriptional activation program and provides resilience against the stress of the beating heart *in vivo*. Mice harboring a mutation in the conserved low-affinity CTCF promoter binding site exhibit an altered, cardiomyocyte-specific transcriptional program and a systolic heart failure phenotype. This transcriptional role for CTCF reveals that a covalent dbTF modification regulating signal-dependent transcription serves as a previously unsuspected component of the oxidative stress response.

## INTRODUCTION

While the ability of many dbTFs to also interact with RNAs has been appreciated for >20 years, this is now complemented by the evidence that low-affinity interactions and multimerization might contribute to the formation of so-called transcriptional “condensates”, often in association with RNAs ^1–3^. This is particularly pertinent in light of evidence that enhancer specificity often depends on submaximal recognition motifs ^4–6^. However, it remains unclear whether RNA association with dbTFs can be regulated by diverse signaling pathways to modulate gene transcriptional programs based on low-affinity sites in regulated transcription units. We elected to investigate this using the *SIRT1* gene as a biologically-important transcription unit, where pioneering studies demonstrated that *Sir2* (Silent information regulator 2) overexpression increases the lifespan of *S. cerevisiae*, whereas loss of function of Sir2 leads to defective epigenetic silencing, deficiencies in DNA repair processes, and shorter lifespan ^7^. These findings were corroborated in other eukaryotes, including *C. elegans* and Drosophila, indicating highly conserved roles for Sir2 ^8–10^. SIRT1, the closest homolog of seven yeast Sir2 homologs in mammals (i.e., SIRT1-SIRT7)^11^, is located in the nucleus ^12,13^, where it catalyzes nicotinamide adenine dinucleotide (NAD+)-dependent histone and non-histone lysine deacetylation^14–16^ to regulate multiple metabolic functions and signaling pathways that link SIRT1 to the normal physiology of various tissues as well as longevity in animal models and in humans ^17–19^. SIRT1 is regarded as a potential mediator of age-related cardiovascular diseases, and its levels decrease with age ^20–22^. SIRT1 plays an active role in the cellular defense against oxidative stress, and its function is affected by the presence of ROS through post-translational modifications ^23–25^. SIRT1 deacetylates FOXO3 and p53 to promote cell survival ^26–29^.

Therefore, the regulation of *Sirtuin 1* (*SIRT1*) transcription by the well-studied chromosome architectural protein CTCF provides a suitable biologically-important model to investigate the largely overlooked potential role of CTCF in signal-induced gene transcription. Although knockdown or induced degradation of CTCF has been suggested to have no clear impact on transcription ^30–33^, recent data suggest that CTCF can modify specific transcriptional events and might be required for regulated transcription at some promoters ^34–38^. These seemingly discrepant observations raise the possibility that CTCF regulation of specific transcriptional programs has gone unnoticed because this CTCF-dependent strategy is primarily triggered in response to specific signals and therefore not readily detectable under basal conditions. Indeed, investigation of *SIRT1* transcriptional regulation has provided a model to dissect the signal-dependent modulation of DNA binding and protein:protein interactions dbTF function and to determine its potential impact on low-affinity regulatory elements in signal-regulated promotors and enhancers. Our study reveals the transcriptional regulation of *SIRT1* serves as a cardioprotective mechanism, based on effective binding of CTCF to the *SIRT1* promoter requiring its stress-induced phosphorylation, to regulate CTCF activity on gene transcription.

## RESULTS

### *SIRT1 SNPs* potentially associated with myocardial infarction (MI)

To gain initial insights into *SIRT1* gene transcription as a model for new principles of promoter/enhancer grammar, we elected to take advantage of the impact of genetic variation in the development of age-dependent health conditions ^39^, including cardiovascular disease (CVD) ^40^. Multiple studies have reported SNPs in *SIRT1* non-coding regions that increase the risk of cardiovascular diseases ^41–43^, including murine model studies emphasizing cardioprotective effects of SIRT1 ^25,44–47^. Many of the known risk factors for CVD, such as hypertension, diabetes mellitus and cigarette smoking, are associated with an elevated production of reactive oxygen species (ROS) ^48^, and cardiomyocytes become increasingly susceptible to ROS in older individuals ^49^.

In concert with evidence that *SIRT1* expression is increased by cellular stress, including oxidative damage or nutrient deprivation *in vitro* ^50^ and *in vivo* ^51^, and the observation that induction of *SIRT1* confers stress resistance in different cell types ^47^, we observed ∼2-fold induction of *SIRT1* mRNA by oxidative stress in several cell culture systems upon treatment with H_2_O_2_ and other DNA damage agents (**Fig. 1a, Fig. S1a**), consistent with the reported capacity of *SIRT1* gene expression to be regulated ^52^. SIRT1 protein level was similarly increased in response to H_2_O_2_ treatment (**Fig. 1a**). As initial evidence of potential transcriptional regulation of *SIRT1*, a small, systematic multidisciplinary study of 83 candidate genes acting in genome maintenance pathways for the San Antonio Longitudinal Study of Aging (SALSA) was conducted using a cohort consisting of 356 healthy Mexican Americans, including 153 old and 203 young individuals, whose aging-associated disease variables were well characterized ^53,54^. In this association analysis of 1,227 tag SNPs for age-related diseases, *SIRT1* was strongly associated with myocardial infarction (MI) as shown in 18 out of 20 tag SNPs (**Suppl. Table 1**). Another cohort, the San Antonio Family Heart Study (SAFHS) that was established to identify genetic risk factors for CVD in Mexican Americans ^55^, was employed. The GWAS for 980 individuals in this cohort showed an association with CVD for 2 of 5 Tag SNPs around the *SIRT1* gene (**Suppl. Table 2**), consistent with its potential association with MI in SALSA. Cohort size in the San Antonio study precludes any statistically clear causal conclusions. We employed these data to identify potential functional variants by analyzing sequences of exons, exon-intron junctions, and regulatory regions, including UTRs and the 4kb promoter region, of the *SIRT1* gene. Sequencing of 14 MI individuals versus 15 controls identified a total of 38 SNPs (**Suppl. Table 3**). The association is enriched in the *SIRT1* promoter, licensing speculation that potential functional variants may reside within this region. Some of the motifs overlap the SNP at the −92 position (rs3740053) in the *SIRT1* promoter and thus might potentially be functionally relevant (**Suppl. Table 4**). Testing the potential importance of candidate promoter SNPs by luciferase reporter gene assays with a ∼1.5kb promoter fragment of the *SIRT1* promoter, we found that the wild-type *SIRT1* promoter associated with SNP rs3740053 was activated after H_2_O_2_ treatment, while the −92A→G MI-risk variant *SIRT1* promoter was not (**Fig. 1b**). None of the other SNPs tested had a significant effect on reporter gene activation (**Fig. 1b**). We confirmed these results by generating HEK293 cells homozygous for the −92G MI risk allele via CRISPR/Cas9 genome editing. In a pair of isogenic lines, HEK293 cells harboring the −92 A→G alleles failed to exhibit H_2_O_2_-induced activation of SIRT1 (**Fig. 1c**) while *SIRT1* was induced in response to H_2_O_2_ in the isogenic wild-type and in response to other genotoxic stress (**Fig. S1b**). Knockdown of CTCF with siRNA inhibited this induction in response to H_2_O_2_ (**Fig. 1d, S1c**). To extend this study, we generated lymphoblastic cell lines (LCLs) harboring the −92G/G haplotype, which were prepared from SALSA individuals homozygous for rs3740053. These alleles co-exist with the other 4 SNPs of the MI-associated common haplotypes. We observed H_2_O_2_-induced binding of CTCF to the wild-type, but not the −92G *SIRT1* promoter (**Fig. 1e, 1f**). ChIP-Seq analysis of wild type HEK293 cells revealed ∼300 promoters gained CTCF binding following treatment with H_2_O_2_ (**Fig. 1g**). GRO-seq data of wild type (−92A/A) vs. (−92G/G) LCL cells also showed that *SIRT1* transcription was induced upon H_2_O_2_ treatment in the former, but not in −92G allele-bearing LCLs, based on promoter pause release (**Fig. 1h**). We selected three up-regulated genes recorded by GRO-seq (*CRNDE*, Chr 16; *RPL37*, Chr 5; and *UBXN4*, Chr 2) to confirm that there was indeed an H_2_O_2_-dependent cohort of induced genes that exhibited increased CTCF promoter binding, and quantitated induced binding of CTCF to their promoters upon H_2_O_2_ treatment, using ChIP-qPCR (**Fig. S1d**).

**Figure 1.**
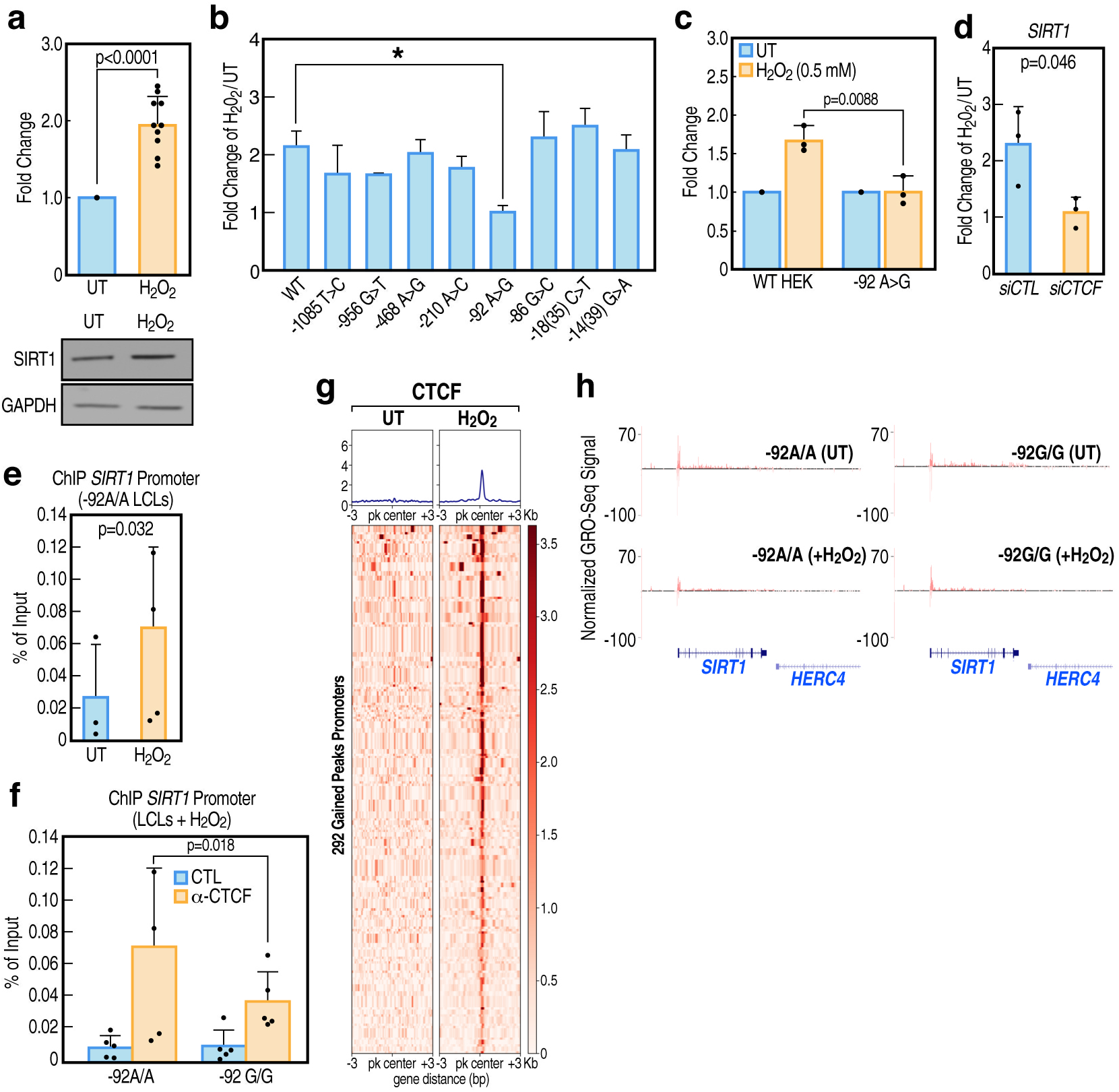
SNP rs3740053 in the SIRT1 promoter impedes oxidative stress induced CTCF regulation of SIRT1 induction. (a) *SIRT1* mRNA levels measured by qPCR and SIRT1 protein expression by Western Blot was increased by H_2_O_2_ in HEK293 cells (15 min 0.5 mM H_2_O_2_, 16 h wash-off). Fold change of *SIRT1* normalized to GAPDH expression relative to untreated (UT) as shown (n=12). (b) A luciferase reporter gene assay containing a 1455 bp promoter fragment upstream of the *SIRT1* TSS was used to compare 24 h wash-off to untreated (UT) cells. The indicated SNP-carrying promoter or wild-type SIRT1 promoter fragment was used. The bar graph shows the fold induction is comparable to the wild-type for all but the −92A>G promoter variant (n=3). (c) CRISPR/Cas9 engineered HEK293 cells carrying the −92A>G mutation (rs3740053) show reduced activation of SIRT1 expression after H_2_O_2_ treatment as measured by qPCR compared to isogenic control HEK293 cells (n=3). (d) Cells transfected with siRNA targeting CTCF show reduced *SIRT1* mRNA induction after H2O2 treatment compared to control siRNA. (e) CTCF ChIP/qPCR shows increased binding of CTCF to the *SIRT1* promoter region including the −92 SNP site in LCL cells treated with 0.25 mM H_2_O_2_ for 15 min, followed by 2 hours of wash-off of H_2_O_2_, compared to untreated control cells. (f) CTCF ChIP-qPCR as in (e) but comparing enrichment of CTCF on the *SIRT1* promoter between LCL cells either homozygous for the A/A majority allele at the −92 position or homozygous for the G/G minority variant. The binding of CTCF is diminished in −92G/G cells upon H_2_O_2_ treatment. (g) CTCF ChIP was performed in HEK293 cells treated for 15 min with 0.25 mM H_2_O_2_ and control cells 2 hours after H_2_O_2_ wash-off. The heatmap shows 292 sites in promoters that show gained binding of CTCF with H_2_O_2_ treatment. (h) Genome browser screenshots of GRO-seq performed in LCLs cells homozygous for either −92A/A or −92G/G SIRT1 after H_2_O_2_ treatment or in untreated cells for comparison. Paused Pol II was detected at both SIRT1 −92A/A and −92G/G genotypes, while H_2_O_2_ treatment leads to pause releases in WT −92 A/A promoters only.

### ROS/Genotoxic stress induces wild-type but not A→G promoter mutant *SIRT1* transcription in cardiomyocytes

To confirm the regulatory events observed for *SIRT1* gene activation in various cell lines apply to events in cardiomyocytes (CMs), we took advantage of the iPSCORE collection of induced pluripotent stem cell-derived cardiomyocytes (iPSC-CMs) ^56,57^, which permitted us to select iPSC-CM lines with either reference or the MI-risk −92A→G variant in the *SIRT1* promoter CTCF site. While the iPSCORE collection has whole genome sequenced lines from 222 ethnically/gender-diverse individuals there were no iPSC-CMs with the homozygous −92G/G genotype, but we identified four lines carrying the heterozygous MI-risk *SIRT1* allele. No significant change in the expression of *SIRT1* under non-stressed conditions was detected between those 4 lines and a set of 16 randomly chosen wild-type iPSC-CM lines **(Fig. 2a)**. We chose two iPSC-CM lines with WT and two iPSC-CM lines from heterozygous MI-risk *SIRT1* alleles to investigate altered transcriptional programs under control versus oxidative stress conditions. We confirmed that the selected cell lines exhibited equivalent basal levels of *SIRT1* expression (**Fig. 2a**) and that *SIRT1* expression increased in response to H_2_O_2_ (**Fig. 2b**). When treated with H_2_O_2_, the reference allele, but not the allele harboring the −92A→G promoter, exhibited activation (**Fig. 2c**).

**Figure 2.**
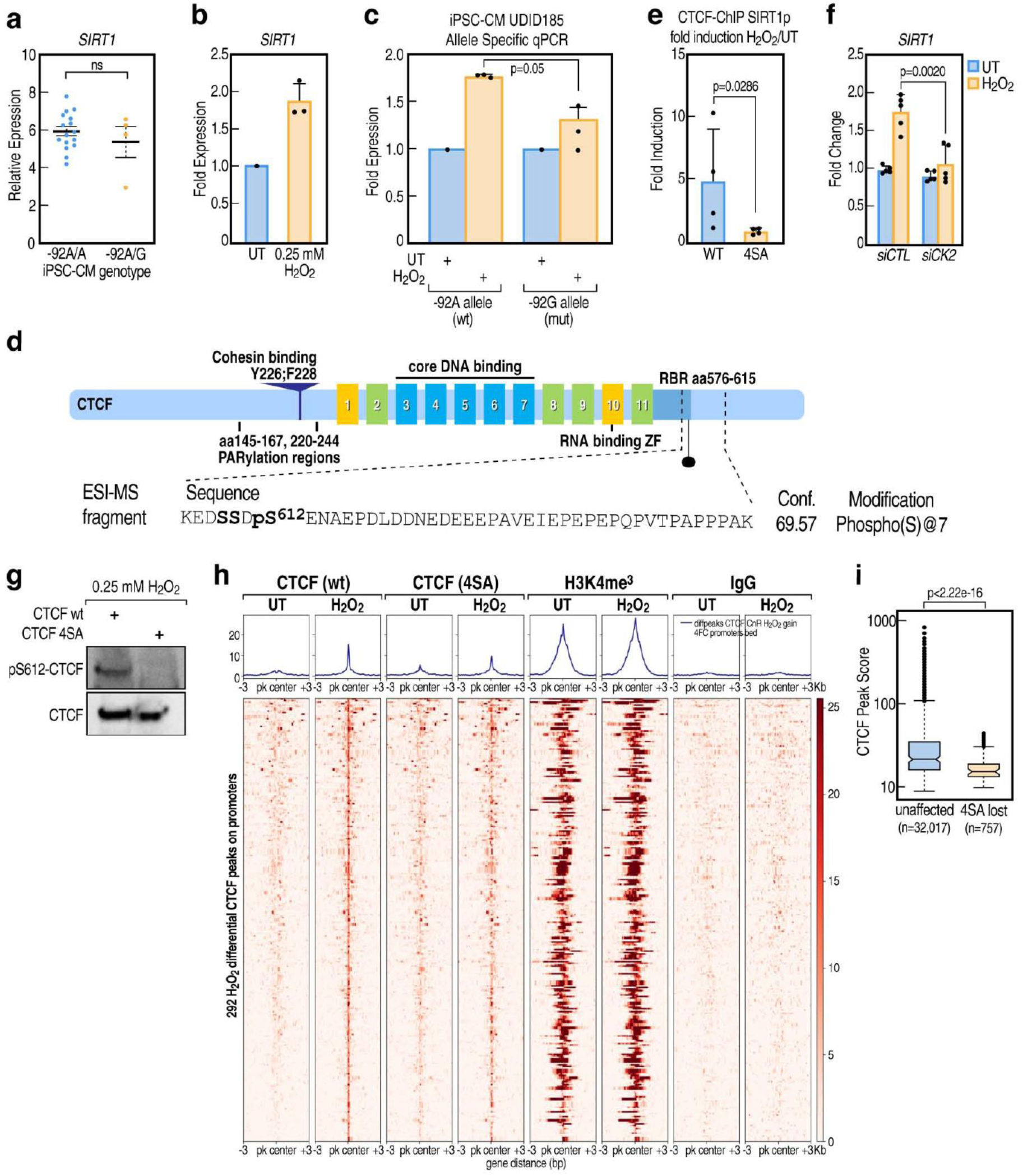
H_2_O_2_-induced phosphorylation of CTCF is a requirement for *SIRT1* regulation in cardiomyocytes and non-cardiomyocyte cells. (a) Relative expression levels of total *SIRT1* are not significantly different between 16 induced pluripotent stem cell-derived-derived cardiomyocytes (iPSC-CMs) with majority (−92A/A) and four with heterozygous (−92A/G) genotype for SNP rs3740053 from the iPSCORE collection. qPCR shows the relative expression of the SIRT1 gene. (b) *SIRT1* mRNA levels in iPSC-CMs measured by qPCR and are increased by H_2_O_2_ (15 min 0.25 mM H_2_O_2_, 8 h wash-off). Fold change of *SIRT1* normalized to GAPDH expression relative to untreated cells (UT) is shown; n=3; p-value = Student’s t-test. (c) Allele-specific qPCR shows induction of *SIRT1* with heterozygous (−92A/G) genotype for SNP rs3740053 (UDID185) after treatment with 0.25 mM H_2_O_2_ for 15 min and samples were processed after 8 h after treatment. Increase of SIRT1 expression after H_2_O_2_ from the −92A allele is significantly more than that from the −92G allele (n=3; p-value = Student’s t-test). (d) Diagrammatic illustration of CTCF protein showing the different Zinc finger (ZF) domains and the phosphorylation site at S612 which was identified by ESI-MS of immunoprecipitated CTCF cells treated with 0.25 mM H_2_O_2_ for 15 min with respect to untreated. S612 resides in the C-terminus of CTCF specifically phosphorylated upon H_2_O_2_ treatment. Detailed results of this assay were tabulated in **Suppl. Table 5**. (e) Endogenous CTCF of HEK293 cells was depleted by 3’-UTR targeting shRNA and reconstituted with Flag-tagged CTCF-WT or CK2-phosphorylation incompetent CTCF-4SA (S604,609,610,612A). Fold-induction of CTCF binding at the at the SIRT1 promoter measured by ChIP-qPCR shows only CTCF-WT, but not CTCF-4SA, that was recruited to the *SIRT1* promoter upon H_2_O_2_ stimulation with 0.25 mM H_2_O_2_ relative to untreated levels (n=4; p-value = Mann-Whitney U-test). (f) Cells transfected with siRNA targeting CK2 catalytic subunit *CSNK2A1* (siCK2) show abrogated *SIRT1* induction after H_2_O_2_ treatment (30 min 0.5 mM H_2_O_2_, 16 h wash-off) compared to control siRNA. (g) Antibodies raised against a peptide with phosphorylated pS612 shows immunoblot specifically detects the phosphorylation of the CTCF S612 residue upon H_2_O_2_ treatment (0.5 mM H_2_O_2_ 15 min, 2 hours of wash-off) in HEK293 cells overexpressing wild-type CTCF but not in the CTCF-4SA mutant. (h) Endogenous CTCF was depleted with siRNA targeting *CTCF* and overexpressing CTCF-WT or CTCF-4SA and treated with H_2_O_2_, analogous to (Fig. 2e). Heatmap of CTCF CUT&RUN shows binding of CTCF in 292 promoters is prevented by mutation of the CK2-phosphorylated serine residues in CTCF-4SA. H3K4me3 and IgG are shown for control. (i) Peak Score of CTCF peaks from CTCF CUT&RUN (Fig. 2h) shows significantly different tag density under 32,017 genome-wide CTCF peaks that unaffected by CK2-phosphorylation mutation CTCF-4SA, compared to 757 peaks that were induced in strength by H_2_O_2_ treatment, but show decreased CTCF binding as an effect of the CTCF-4SA mutation. Bar graphs show the mean +/-sd and the p-value was calculated using students’ two-tailed t-test (*p<0.05; **p<0.01; *** p=<0.01), unless otherwise specified.

### Requirement of H_2_O_2_-induced phosphorylation of CTCF

To evaluate potential posttranscriptional modifications of CTCF upon H_2_O_2_ treatment that might be responsible for its activator function on the *SIRT1* gene expression, we noted that phosphorylation of S612 has been reported to regulate CTCF activity ^58,59^. To confirm that this posttranslational modification occurred in response to H_2_O_2,_ we transiently transfected pCMV10-3xFlag-CTCF plasmid into HEK293 cells, treated with H_2_O_2_ 30 h post-transfection, collected cells at 48 h post-transfection, and following IP with anti-Flag antibody the eluted proteins were digested and analyzed by mass spectrometry (**Suppl. Table 5**). These experiments identified H_2_O_2_-dependent phosphorylation on S612 located in the intrinsically disordered C-terminus of CTCF encompassing 3 consensus sites for protein kinase CK2-mediated phosphorylation at S609, S610, S612, and a possible 4^th^ CK2 consensus site at S604 (**Fig. 2d, Suppl. Table 5**). Because ROS/H_2_O_2_ activates Ataxia telangiectasia and Rad3-related (ATR) kinase ^60^ and CK2 ^61^ we performed knockdown of endogenous CTCF employing 3’-UTR-specific shRNAs and reconstitution with either Flag-tagged wild-type (WT) or mutant CTCF (S604,S609,S610,S612→A, referred to as 4SA), where the four adjacent CK2-phosphorylated serine residues are replaced by alanine. Anti-Flag ChIP experiments showed that only reconstitution with Flag-CTCF-WT but not with Flag-CTCF-4SA, restored recruitment of CTCF at the SIRT1 promoter after treatment with H_2_O_2_ (**Fig. 2e**). Mutation of the serine residues to alanine eliminated the ability of H_2_O_2_ to increase CTCF binding to the *SIRT1* promoter or to activate *SIRT1* gene expression (**Fig. 2e, S2d**). Consistent with this observation, knockdown of the gene encoding CK2 catalytic subunit, *CSNK2A1*, abolished *SIRT*1 induction (**Fig. 2f**). To provide additional evidence, we raised and purified anti-phospho-peptide IgG corresponding to this C-terminal region of CTCF and detected CTCF phosphorylation in response to H_2_O_2_ by Western blot analysis, finding that this antibody bound to WT CTCF, but did not bind to CTCF harboring a S612A mutation of the most robust CK2 phosphorylation site following H_2_O_2_ treatment (**Fig. 2g**). Unfortunately, this antibody proved insufficiently sensitive for use in either ChIP-seq or CUT&RUN assays.

To obtain a global perspective on the impact of CTCF phosphorylation site mutants on signal-dependent recruitment of CTCF, we performed CUT&RUN assays in HEK293 cells expressing either the WT or 4SA CTCF mutant. This analysis revealed that 90-300 promoters (depending on cut-off) gained low levels of CTCF binding in response to H_2_O_2_ treatment, which was not observed for the 4SA phosphorylation-defective CTCF mutant protein (**Fig. 2h, S2e**). These data also indicate that the strength of CTCF binding at the gained promoters is only ∼1/4-1/10th that at the strong CTCF binding sites, such as those at TAD boundaries (**Fig. 2i**). In contrast, strong CTCF sites, such as those at TAD boundaries, did not show clear alterations in binding in response to H_2_O_2_ (**Fig. S2b**). Total CTCF peaks and induced CTCF promoter binding in response to H_2_O_2_ treatment decreased (**Fig. 2i**).

### H_2_O_2_-induced Architectural Changes License *SIRT1* Gene Activation

We next applied 4C to determine whether the *SIRT1* promoter exhibited induced interactions with a strong CTCF site located ∼30kb 5’ of the *SIRT1* promoter. Using the −30kb site as the viewpoint, these experiments revealed that treatment with H_2_O_2_ elicited an interaction of the *SIRT1* promoter with the −30kb strong CTCF binding region in WT HEK293 cells. This was not observed in HEK293 cells homozygous for the −92A→G mutation (**Fig. 3a**). Because of the induced interaction and hence the altered architecture of the *SIRT1* gene locus, we next explored the potential role of the lncRNA at −35kb. A browser image of the *SIRT1* locus shows the transcription of the −35kb lncRNA/enhancer, the induction of the *SIRT1* locus, and the binding of CTCF to both a strong −30kb site and to the weak *SIRT1* promoter site in response to H_2_O_2_ (**Fig. 3b**).

**Figure 3.**
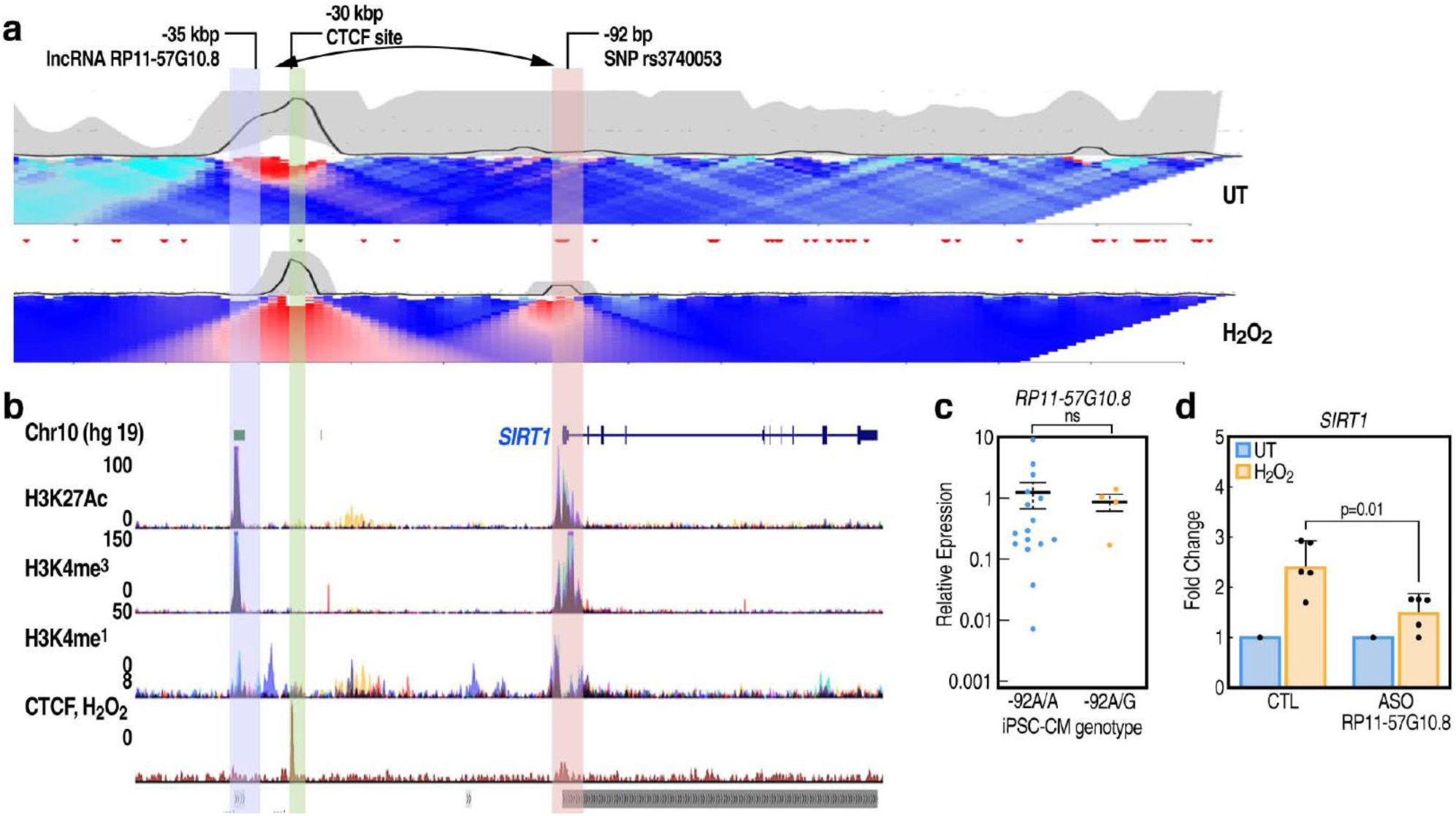
Oxidative stres*s* regulates DNA looping of SIRT1 promoter to lncRNA RP11-57G10.8. (a) 4C-seq map of untreated and H_2_O_2_ treated (15 min 0.25 mM H_2_O_2_, 1 h wash-off) in HEK293 cells. The viewpoint around the −30kb CTCF site shows increased interaction with the *SIRT1* promoter region after H_2_O_2_ treatment. (b) CTCF ChIP (bottom row) showing the binding strength of CTCF proteins at the SIRT1 promoter is a fraction of that on the −30kbp site. ENCODE genome browser tracks for H3K27Ac, H3K4me3, and H3K4me1 show deposition of those histone marks at the *SIRT1* locus in multiple cell lines. (c) Relative expression levels of *RP11-57G10.*8 lncRNA in a total of 16 induced pluripotent stem cell-derived-derived Cardiomyocytes (iPSC-CMS) with wild-type and four with heterozygous genotype for SNP rs3740053. qPCR shows the relative expression of the lncRNA and GAPDH was used as a normalization control. (d) Relative fold change expression of SIRT1 in either control ASO or ASOs against *RP11-57G10.8*-transfected HEK293 cells were treated with 0.5 mM H_2_O_2_ or left UT for 1 hour followed by RNA isolation after 24 hours of treatment and *SIRT1*expression was measured by qPCR.

Previous studies have described CTCF dimerization and multimerization ^62–64^, noting that multimerization was impaired by RNase treatment ^64–67^. Further, this RNA-modulated CTCF multimerization was variously reported to be inhibited by deletion of zinc finger ZF1, ZF10, ^65^ and a C-terminal RNA-binding region of CTCF 3’ of ZF11 (aa576-615, see **Fig. 2d**), that has also been implicated in RNA-facilitated CTCF multimerization ^66^. Therefore, we assessed whether lncRNA *RP11-57G10.8,* located −35kb from the *SIRT1* promoter, might prove to be the RNA moiety actually required to augment the multimerization of phosphorylated CTCF to license stable binding to the *SIRT1* promoter. The levels of lncRNA *RP11-57G10.8* were essentially unaltered in cells with either WT or −92A/G *SIRT1* genotype (**Fig. 3c**). ASO treatment to reduce *RP11-57G10.8* levels proved sufficient to reduce induction of the *SIRT1* promoter by ∼75% in response to genotoxic stress (**Fig. 3d**), revealing the functional requirement of this adjacent lncRNA for effective *SIRT1* transcriptional activation.

### *In vivo* evidence of the role of induced CTCF binding on *Sirt1* promoter function

Having established that stress-induced increased transcription of *SIRT1* occurs in cardiomyocytes and that this is based on phosphorylated CTCF, licensing its binding to the low affinity *SIRT1* promoter CTCF site, it was important to determine whether this low affinity CTCF site would be important for *SIRT1* activation *in vivo* and whether the 2-fold transcriptional activation was of clear biological significance. Indeed, a low affinity CTCF binding site is a conserved feature of the *SIRT1* promoter in multiple species, including mouse (**Fig. 4a**); however, aside from the core region there is sequence variation between species. Employing a CRISPR/Cas9 strategy to generate appropriate *Sirt1* promoter CTCF site mutants (Sirt1-Mut) mice, we were able to create two founder lines that had the same two base pair CTCF motif mutation in the *Sirt1* promoter (**Fig. 4a**). Continued breeding of the Sirt1-Mut lines exhibited no discrepancies in the expected Mendelian ratios during genotyping. Homozygous Sirt1-Mut mice of both sexes had a normal developmental progression, showed normal behavior and aging as their wild-type littermates and heterozygous animals.

**Figure 4.**
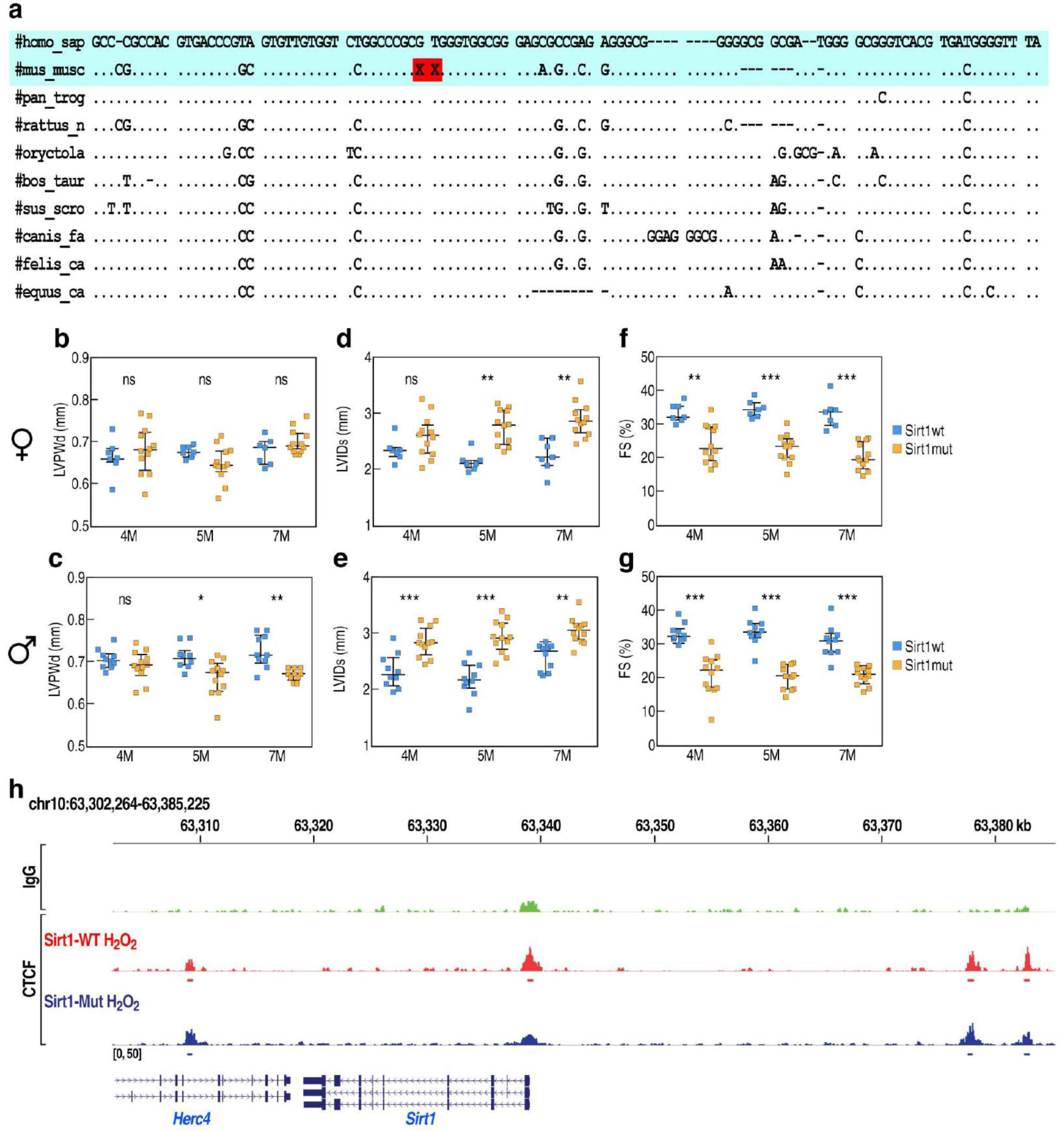
Echocardiography and gene expression patterns in hearts of wild-type and Sirt1-promoter mutant mice. (a) *SIRT1* promoter low affinity CTCF site conservation in various species human, mouse, chimpanzee, rat, rabbit cattle, pig, dog, cat, horse (top to bottom). “X” represents the site of deletion in Sirt1-Mut mice. (**b,d,f**) Female (n=12 Sirt1-Mut, orange; n=7 WT littermate control, blue) and (**c,e,g**) male (n=12 Sirt1-Mut, orange; n=10 WT littermate control, blue) mice at ages 4, 5 and 7 month (4M, 5M, 7M). Left ventricle posterior wall thickness; end diastole (LVPWd, mm); left ventricular internal diameter, end systole (LVIDs, mm), Fractional Shortening (FS, %) data from echocardiography; 2-way ANOVA, Bonferroni correction p = * < .05, ** < .01; *** < .001. (**h**) Browser data of the CTCF CUT&RUN data of neonatal mouse ventricular myocytes (NMVM) derived from wild-type (SIRT1-WT) and Sirt1-Mut mice treated with H_2_O_2_ for 15 min, 2 hours recovery.

To directly test whether the CTCF binding site mutation in the *Sirt1* promoter influences cardiac function, we had initially anticipated a requirement for subjecting wild-type and Sirt1-Mut littermates to Traverse Aortic Constriction (TAC) or sham operation. However, echocardiography performed to assess baseline cardiac health on a cohort of 7-12 age-matched 4-month-old mice revealed an unanticipated phenotype in the mutant mice. 2D echocardiograms were analyzed to record systolic and diastolic cardiac function, wall thicknesses inner cardiac dimensions, morphology, and mass (**Suppl. Table 6**). The body weight of animals was not significantly different between *Sirt1-promoter mutant* and littermate groups over time, making it unlikely that the results were influenced by differences in body size (**Suppl. Table 6**). Echocardiographic images (two-dimensional B-Mode, M-mode, tissue Doppler, and trans-mitral pulse-wave Doppler) were recorded and QRS-guided echocardiographic parameters including diastolic interventricular septum thickness (IVSd), systolic and diastolic left ventricular internal diameter (LVIDd, LVIDs), fractional shortening (FS) (**Fig. 4b-g**) were measured. Indeed, for cohort of male *Sirt1* promoter-mutant mice, we observed that at 4 months, and in the absence of exogenous stress, echocardiography revealed LV hypertrophy in these mice and evidence of cardiac insufficiency, i.e., show reduced FS, and increased LVIDs wall dimensions, which were not observed in littermate control mice (**Fig. 4c, 4e, 4g**). Therefore, this specific mutation in the *Sirt1* promoter CTCF site was sufficient, even in the absence of overt exogenous cardiac stress, to initiate cardiac hypertrophy/failure in young male mice.

Successive echocardiography in 5- and 7-month-old male mice showed mildly progressive heart failure in Sirt1-Mut mice. 7-month-old male mice Sirt1-Mut mice show reduced FS% (control 31.1 ± 1.5%, Sirt1-Mut 20.9 ± 0.8%, mean ± sem, p < 0.0001, n=10-12 each), as well as increased LVIDs and decreased left ventricle posterior wall (LVPWd) dimensions (**Figure 4c, 4e, 4g**). A similar type of cardiac failure was observed in female mice, but as expected, was milder than that observed in male mice (**Figure 4b, 4d, 4f**), with analysis of mice at 4, 7 and 9 months of age revealing a mild progression of the pathology over time both in male and female mice.

To ascertain the potential actions of CTCF on the *Sirt1* promoter in the myocardium, we prepared cardiomyocytes from neonatal WT or Sirt1-Mut mice and performed CUT&RUN analyses. Consistent with the effects of genotoxic stress on expression of *SIRT1*, we noted that the level of CTCF binding on the *Sirt1* promoter is observed in WT mice but not in homozygous *Sirt1-* mutant mice littermates (**Fig. 4h**), consistent with the importance of up-regulation of in response the stress of the physiologically beating heart.

### Sirt1-Mut mice exhibit a cardiomyocyte-specific altered transcriptional program

To examine whether the altered function reflected actions in cardiomyocytes, or in additional cell types, we performed bulk and snRNA-seq analysis of left ventricles (LV) of wild-type and Sirt1-Mut male mice, with sufficient cell representation to definitively analyze cardiomyocytes, cardiac fibroblasts, and endothelium (**Fig. 5a, 5b**). For bulk analysis, 8-month-old Sirt1-Mut mice and control littermates were sacrificed and LV biopsies were collected for RNA extraction. RNA-seq of LV biopsies revealed a program of >120 differentially expressed transcription units in the myocardial tissue (**Fig. 5a**). This is estimated as 55 up-regulated gene transcripts, with Gene Ontology (GO) term analysis revealing regulation of oxygen species, regulation of oxidative stress-induced cell death among the top changed terms (**Fig. S3a**). 75 transcription units were down regulated, with GO term analysis showing the regulation of chemokine secretion and dendritic cell chemotaxis were affected (**Fig. 5a, S3a**). We next performed single nucleus RNA-seq on cells collected from the hearts of two 9-month-old Sirt1-Mut mice and two wild type littermates to assess transcriptional differences between WT and Sirt1-Mut individuals. Analysis of differentially expressed genes in WT vs Sirt1-Mut tissue confirmed that mutation of the *Sirt1* promoter CTCF site resulted in decreased *SIRT1* transcripts in cardiomyocytes (**Fig. 5c**). Comparing individual mice material by snRNA-seq, we observed a similar but not identical pattern of transcripts exhibiting up and down-regulation in mice with the mutated promoter CTCF site, the vast majority of which are expressed exclusively in cardiomyocytes (**Fig. 5b, 5d**). Indeed, the *Sirt1* transcript was decreased in cardiomyocytes in male Sirt1-Mut mice compared to WT mice (**Fig. 5c**). Volcano plots of up- or down-regulated transcripts in Sirt1-Mut myocardial tissue by snRNA-seq analysis revealed the larger number of down-regulated vs up-regulated transcripts in male Sirt1-Mut cardiomyocytes, with only a small subset of altered transcripts observed in fibroblasts and almost no altered transcripts in endothelial cells (**Fig. 5d, S3b,c**). Examination of the GO terms for biological process the up- and down-regulated transcriptional profile, and their statistical significance, is summarized in (**Fig. 5e)**. These data are consistent with the observation that >85% of the genes altered in bulk analysis occurred in cardiomyocytes (**Fig. S3c**), with the prominent GO terms being abnormal cardiac fiber morphology, cardiac hypertrophy, and regulation of relaxation of cardiac muscle (**Fig. S3d**).

**Figure 5.**
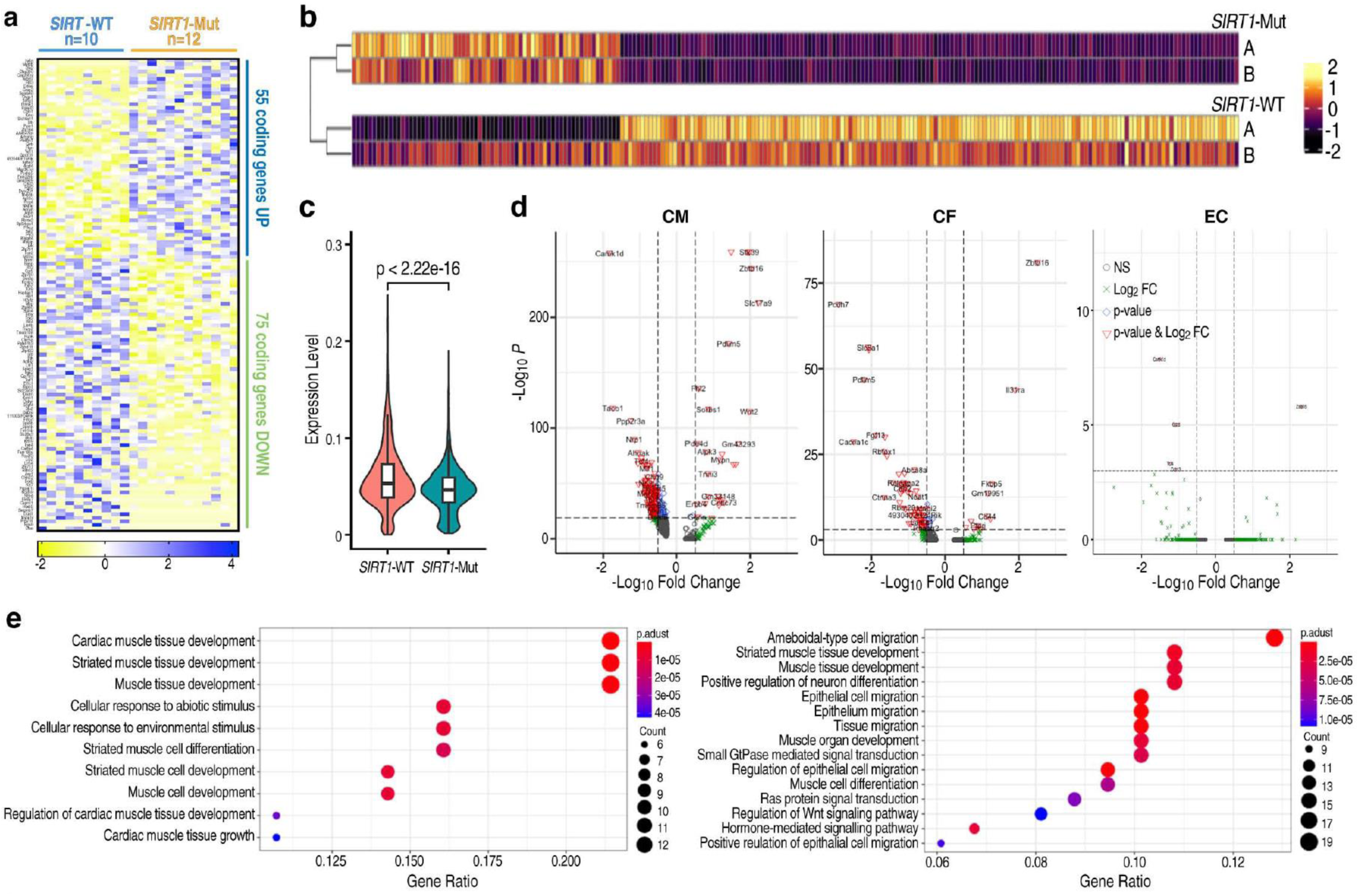
snRNA-seq shows an altered transcriptional program in cardiomyocytes of Sirt1-Mut mice. (a) Bulk RNA-seq was performed on left ventricle (LV) apex biopsies of 12 male Sirt1-Mut mice and 10 wild-type littermates (SIRT1-WT), showing differential expression resulting in the upregulation of 55 genes and 75 downregulated protein-coding genes, as shown in heatmap. (b) snRNA-seq was performed in duplicate on two male 9-month-old Sirt1-Mut mice and two wild-type littermates, respectively, resulting in similar but not identical differentially expressed genes, as shown in heatmap. (c) snRNA-seq reflecting the lower level of SIRT1 expression in Sirt1-Mut mice compared to WT mice. Implicating that presence of SNP in SIRT1 promoter results in lower SIRT1 expression in mice as compared to WT. (d) Volcano plots showing the differentially expressed genes identified from snRNA-seq of wild-type control or Sirt1-Mut male mice cardiomyocytes (CM), cardio fibroblasts (CF), and endocardial cells (EC). Cardiomyocytes show the predominant proportion of differentially expressed genes. (e) Gene Ontology (GO) terms analysis of differentially expressed (down-, left panel; and up-regulated, right panel, respectively) genes in snRNA-seq cardiomyocytes representing defects in cardiac fiber morphology and relaxation of cardiac muscles.

These data reveal that a *CTCF* binding site mutation in the murine *Sirt1* promoter, critical for oxidative stress responses, altering *Sirt1* response to stress, leads to cardiac dysfunction, featuring defective systolic relaxation (**Fig. S4**). This occurs in mice without any experimental intervention to induce conditions mimicking human heart failure. The transcriptional regulation of *Sirt1* in response to stress thus appears to play an important, largely unappreciated, cardioprotective function. These results are consistent with reports that *SIRT1* can induce nuclear factor erythroid 2-related factor 2 (Nrf2) ^68–70^ in endothelial cells and cardiomyocytes ^71^, where it triggers anti-apoptotic functions ^46^, counteracts endoplasmic reticulum stress ^72^, increases myocardial contractility ^73^ and facilitates resistance to ischemia/reperfusion injury ^45^.

## CONCLUSIONS

Many DNA-binding transcription factors (dbTFs), as well as architectural chromatin-associated proteins such as CTCF, engage in functional low-affinity interactions which becomes particularly pertinent with evidence that physiological enhancer activation programs often depend on submaximal recognition motifs ^4,5^ a principle whereby even single nucleotide variants within heart enhancers can disrupt heart development ^6^. Here, investigating this issue we present evidence establishing that posttranscriptional modification of CTCF after oxidative stress can critically modulate DNA binding with low-affinity sites. Both *in vitro* and *in vivo* data point to the prevalence of low affinity sites in the human genome ^74^, and regulatory activity is a feature of low affinity sites during development that can be disturbed by single nucleotide variants ^75^. Thus, we were able to identify a SNP that even further lowers the CTCF binding affinity on *SIRT1* promoter site abolishes its ability to respond to genotoxic stress (**Fig. S4)**. On the other hand, mutations of CTCF that impair multimerization, such as mutations in zinc finger 10, similarly preclude effective promoter binding to the low-affinity *SIRT1* promoter site, again abolishing stress-induced activation. However, this is required for binding to the low affinity promoter sites in stress-regulated transcription units. Critically, stress-induced phosphorylation of CTCF is necessary for binding to the low-affinity promoter DNA sites. Given the importance of low affinity dbTF binding sites throughout the genome, we suggest that signal-regulated binding will be an important general regulatory strategy for promoter and enhancer grammar.

Our data reveal that acute signal (H_2_O_2_)-dependent stimulation causes specific C-terminal phosphorylation of CTCF and binding to low-affinity DNA promoter sites, resulting in transcriptional activation of the *SIRT1* gene. We note that, in response to genotoxic stress, increased CTCF binding can additionally be detected on many low affinity sites, and that this is accompanied by activation of these coding transcription units. *In vivo* experiments reveal that SIRT1 transcriptional activation protects against a specific form of cardiac failure, marked by LV hypertrophy and evidence of cardiac insufficiency, with reduced fractional shortening, and increased left ventricular internal diameter wall dimensions, indicating that this mechanism serves as a component of the resilience program that protects against the normal physiological stress of the beating heart. Single nucleus RNA-seq analyses indicate that the observed failure of systolic relaxation involves an altered cardiomyocyte-specific transcriptional program.

This study provides an addendum to the reports that knockdown or proteolytic degradation of CTCF using the degron system exerts minimal or no effects on gene transcription under basal conditions ^31,32,76,77^, revealing a signal-dependent transcriptional role for CTCF. CTCF is recruited to low-affinity sites on a cohort of promoters, including *SIRT1,* in response to CK2-dependent phosphorylation of specific C-terminal serine residues that are adjacent to regions of CTCF suggested to be involved in RNA interactions and mediate RNA-dependent CTCF multimerization ^65,66^. These conclusions are in concert with findings of CTCF-dependent interactions with some promoters ^78^ and genetic data revealing that promoter binding of CTCF can be functionally required for ligand-dependent enhancer:promoter interactions ^79–81^. This study reveals that covalent modification of CTCF can serve as a previously unsuspected strategy to regulating signal-dependent DNA binding, thereby serving as a broadly-used component of promoter grammar regulation in metazoans.

## Contributions

The study was conceived and initiated by YS and MGR, with the initial validating experiments by XS and MC. SALSA and SAHFS data analysis by SY, in collaboration with HH, RD, JB. The data for manuscript figures for the manuscript were primarily contributed by TW, PP, RM and MF, with help from MA. Mass spectrometry experiments were prepared by SJN. CTCF antibody and testing was implemented by TW. Lymphoblastoid lines were prepared by SD, iPSC-CMs were prepared AD’A-C, MD’A, and KAF. The mouse model was established and maintained by TW and HT, and the echocardiography, tissue harvesting, and cardiac RNA extraction was performed by AEZ-H, RM and HT, who managed the colony. snRNA-seq was performed by RM and TW. CG, RM and AG performed bioinformatic analyses, TW, PP and RM performed CUT&RUN experiments; XS, SN, PP and TW performed ChIP and ChIP-seq experiments; TW and SO performed 4C experiments. SJN, KAO, TW, PP, MA and MJF prepared the CTCF mutants. RM, TW and KAO prepared sequencing libraries. TW, PP, YS and MGR wrote the manuscript, with particular input and discussion from AKA, AEZ-H and input from all authors.

## Acknowledgements

We thank Jun Zhao in the UCSD transgenic core for preparation of the Sirt1-Mut mouse lines, Kristen Jepsen and the UCSD IGM core for sequencing, and Majid Ghassemian from the UCSD Biomolecular and Proteomics Mass Spectrometry Facility. We thank Janet Hightower for help with figure preparation. This publication includes data generated at the UC San Diego IGM Genomics Center utilizing an Illumina NovaSeq 6000 that was purchased with funding from a National Institutes of Health SIG grant (#S10 OD026929) and from the UCSD Biomolecular and Proteomics Mass Spectrometry Facility utilizing an Thermo Scientific Orbitrap Fusion Lumos (#S10 OD021724). This study was supported by grants from National Heart, Lung, and Blood Institute (R01HL150521 to MGR), National Institute of General Medical Sciences (R35GM150636 to SNJ), and American Herat Association Postdoctoral Fellowship 19POST34450099 (TW).

## MATERIALS AND METHODS

### SNP selection and Genotyping

We selected our candidate genes of which resequencing data is available in Environmental Genome Project (EGP) database (http://egp.gs.washington.edu/) and used tagSNPs listed in the order forms provided by the database for genotyping. These SNPs are chosen by LDSelect, a SNP selection algorithm based on the linkage disequilibrium statistic r². We initially pooled all the SNPs in these order forms of our candidate genes and scored for a customized bead array, Illumina GoldenGate which genotypes 1,536 SNPs in an assay. The initial number of SNPs used for scoring and tested for designability was 2,052 in 89 genes. After final scoring process, 1,384 SNPs in 83 genes were left to be assayed by dropping SNPs with the score lower than 6.0 and designability rank lower than 1. After genotyping and quality control, 1,227 SNPs were used for association analysis.

### SALSA and SAFHS Study Population

The San Antonia Longitudinal Study on Aging (SALSA) tracked the incidence of physical impairments, disability, and handicap, including cardiovascular diseases in a cross-sectional functional status assessment among elderly members (64+ years old) of the San Antonio Heart Study cohort (SAHS) in the San Antonio, Texas, region ^53^.

The San Antonio Family Heart Study (SAFHS) is a longitudinal study with information from 2,590 Mexican American individuals from San Antonio, Texas, designed to identify low frequency or rare variants cardiovascular disease, using whole genome sequence ^82^.

### Cell Culture, transfections and treatment

The protocol for cell culture followed previous work ^79^. In brief, we originally purchased COS-7, HEK293T and HEK293 cells from ATCC, which were maintained in DMEM (Gibco, 10566) supplemented with 10% FBS (FB-11, Omega Scientific) in a 5% CO2 humidified incubator at 37 °C. LCL cells were generated as described previously. Briefly, LCL suspension cells were cultured in RPMI 1640 media (Gibco, 11875093) with 15% FBS. Cells were examined for mycoplasma contamination every 6-12 months. For the four iPSC-derived cardiomyocyte lines (iPSC-CMs; iPSCORE_103_1 “UDID104”; iPSCORE_14_2 “UDID105”; iPSCORE_54_1 “UDID106”; iPSCORE_84_1 “UDID185”) ^57^, briefly, 1M differentiated iPSC-CMs were seeded in a Matrigel-coated (Corning, 354230) 6-well plate in RPMI supplemented with 20% FBS, B-27 Supplement (Gibco, 17504044), and 10 mM ROCK Inhibitor (Sigma, Y0503). Starting 24 hours after plating, the medium was replaced with serum free RPMI supplemented with B-27 Supplement and changed every 2 days for at least 14 days and checked for signs of cardiomyocyte activity (i.e. beating) daily. After initial isolation, NMVMs were weaned off serum by one passage in DMEM with half amount of serum, and then further cultured in serum starved (<1%) DMEM to reduce non-NMVM growth. Medium was changed every day.

Transfections of HEK293 cells with siRNA, ASO and plasmids at 60–80% confluency was performed using Lipofectamine 2000 (Life Technologies) following the manufacturer’s instructions. For transfection of LCL cells with siRNA using Lipofectamine 2000, cells were passaged and a reverse transfection protocol with 1 million cells was used according to the manufacturer’s instructions. The control si-, sh-, and ASO oligos used in this study include Santa Cruz negative control siRNA (sc-36869), IDT control B ASO (QTE-240580), Sigma Mission siRNA universal control #2 (SIC002).

For induction of Reactive Oxidative Species (ROS) stress in tissue culture, cells were treated with 0.25-0.5 mM H2O2 (Sigma, H1009) for 15-30 min. After treatment, the H_2_O_2_ containing medium was removed/washed-off and replaced with fresh medium for further incubation until sample collection (1-2 h for ChIP, Run-On, and CUT&RUN; 6-24 h for RNA). Treatment with etoposide (Sigma, E1383) was continuous until sample collection.

### CRISPR/Cas9 generation of mutant cell lines

For the genotyping of SIRT1 SNPs selected from sequencing results, we used both conventional Sanger sequencing and Sequenom MassARRAY platform.

### Luciferase Assay

The Luciferase reporter is constructed by cloning 1,455bp SIRT1promoter region into pGL2 Basic vector using XhoI and MluI restriction sites. The insert was amplified by 5’-CATGCTC-GAGCTTCCAACTGCCTCTCTGGC-3’ and 5’-CATCACGCGTCGCATGCATTAGCTATT-TGTCCTA-3’from a genomic DNA prepared from one of the subjects in our population who carry major alleles for all the SNP sites using i-Proof High Fidelity Polymerase (Bio-Rad). The individual associated SNPs were introduced by QuikChange® II XL Site-Directed mutagenesis kit (Stratagene, 200523). All clones were confirmed by sequencing. COS-7 cells were transfected with reporter plasmid construct together with pRL-TK Renilla luciferase plasmid by Lipofectamine™ 2000 (Invitrogen).

### qPCR

RNA was isolated using Zymo MiniPrep RNA prep kit (R1055, Zymo Research) with DNase I treatment. The total RNA was reverse-transcribed using SuperScript III Reverse Transcriptase (Life Technologies, 18080-051) with random hexamer and oligo-dT primers as per the manufacturer’s instructions. qPCR was performed on Mx3000P qPCR systems (Agilent) using VeriQuest FastSYBR 2X qPCR master mix (Thermo, C-75690). Normalization of expression was done using GAPDH or ACTB mRNA as internal controls. For all RT-qPCR and ChIP-qPCR, experiments were performed with at least two replicates, and technical duplicates or triplicates for each biological sample. A list of primers used for qPCR is provided in **Suppl. Table 7**. If not indicated otherwise, p values were obtained using a two-tailed Student’s t-test, * = p<0.05 and ** = p<0.01, or as indicated.

### Western Blot, immunoprecipitation (IP)

HEK293 and HEK293T cells co-transfected with HA- and Flag-tagged CTCF were treated with H2O2 were either treated with H_2_O_2_ for 30 min or remained untreated. Samples were harvested in 350 µl of RIPA buffer (Thermo, 89900) supplemented with Protease and phosphatase inhibitor cocktail (Thermo, 78440) and tip-sonicated (Branson sonifier, 10 s) on ice. The lysate was centrifuged at 10000 rpm for 10 min at 4°C and the supernatants were used fresh or stored at −80°. For immunoprecipitation (IP) experiments, the lysate was incubated for 4 hours with 1 ug of anti-HA or anti-Flag antibody at 4°C on a rotator. 1% of the lysate was kept aside as input. Protein-G beads were washed, blocked with 1% BSA in TE buffer, and then added to the lysate and for 1 hour IP at 4°C. Magnetic beads were captured on a magnet and washing with RIPA buffer 3 time rotating for 5 min. 4x Loading buffer (Thermo, NP0007) was added to beads and input was boiled and used for Immunoblotting.

Western blotting was performed as described ^83^ with modifications using XCell II™ Electrophoresis and Blot System (Thermo), using precast Bis-Tris Gels (NP0323BOX, Thermo) and MOPS SDS Running buffer (MB1035, Biopioneer) for SDS-PAGE followed by transfer to Nitrocellulose membranes (1620115, BioRad) in Transfer buffer (MB1037, Biopioneer). Membranes were blocked in 5% BSA or non-fat dry milk in PBS + 0.05% Tween-20, incubated with primary antibodies 4 h to over-night at 4°C and with HRP-coupled secondary antibodies 1 h at room temperature in 2% BSA or NFDM in PBS-Tween. ECL substrates (34580, Thermo) were used for signal detection with X-ray film (MUBF-02, Biopioneer) with an automated film developer () or a chemiluminescent imaging system (PXi, Syngene).

### Mass Spectrometry

Briefly, target enrichment of Flag-CTCF expressed in HEK293T cells treated with 0.25 mM H_2_O_2_ (15 min, 8 hours recovery) and left untreated cells, were enriched by anti-Flag-IP and recovered by gel electrophoresis. The gel was stained and the CTCF band cut out and handed over to the. UCSD Biomolecular and Proteomics Mass Spectrometry Facility for phospho-peptide enrichment (IMAC), large scale in solution digest, LC (2 hour reverse phase C18 gradient)-MS2 mass spectrometry and data analysis to identify posttranslational modifications on CTCF.

### Chromatin Immunoprecipitation (ChIP) and ChIP-seq

ChIP was performed as previously described ref ^79^. In brief, cells were either cross-linked directly with 1% formaldehyde at room temperature for 10 min, or cells were double cross-linked with 1 mM DSG (ProteoChem) for 1 h followed by 10 min 1% formaldehyde. In both situations, the cross-linking was quenched by addition of 0.125M glycine for 10 min. Chromatin was fragmented using sonication (Bioruptor Pico, Diagenode) (10–30 cycles, 30 s on/30 s off). Subsequently, the soluble chromatin was cleared by centrifugation (10000 g, 10 min 4°C), pre-cleared with 10-20 μl Protein G Dynabeads (10009D, Life Technologies), and then incubated with 1–5 μg of antibodies at 4 °C overnight. ChIP complexes were collected using 30 μl of Protein G Dynabeads per sample for the last hour of incubation. The immune complexes were subjected to washes once with wash buffer I, twice with wash buffer II, once with Tris-EDTA (TE) + 0.1% Triton X-100, and once with TE, and then the beads were incubated at 55 °C for 2 h with proteinase K and de-cross-linked at 65 °C overnight. The final ChIP DNA was extracted and purified using QIAquick columns. For ChIP-seq, the extracted DNA was ligated to specific adaptors for Illumina’s HiSeq system using the KAPA Hyper Prep Kit (Kapa Biosystems).

### CUT&RUN-seq

CUT&RUN was performed as previously described ^84^ with a CUTANA™ ChIC/CUT&RUN Kit (14-1048, Epicypher) according to the manufacturer’s instruction. Briefly, cells were harvested, counted, and 0.5 M fresh live cells used per CUT&RUN reaction. Alternatively, nuclei were isolated from 0.5 M cells and stored at −80°C. Cells or nuclei were immobilized on ConA magnetic beads, permeabilized with Digitonin and incubated with specific or control antibodies over-night on a Nutator. After MNase binding and activation with Ca^2+^, the released DNA fragments were isolated from the supernatant. DNA amount was measured by Qubit and NEBNext Ultra II (NEB) used for library prep. Library size distribution was evaluated on Bioanylazer (Agilent) and sequenced on NovaSeq 6000 (PE100).

Sequencing reads were inspected for quality control using FASTQC (https://www.bioinformatics.babraham.ac.uk/projects/fastqc/) and sequencing adaptors were trimmed, if necessary, using TRIMMOMATIC (Version 0.39) (http://www.usadellab.org/cms/?page=trimmomatic). Reads were aligned to hg19 with Bowtie2 ^85^ (version 2.26) using --very-sensitive setting and --dovetail option for overlapping reads. Tag directories were then generated using HOMER ^86^ (version 4.11.1), keeping only unique aligned reads per genome position, allowing one unique read per position (−tbp 1).

CTCF peaks were called using HOMER ^86^ findPeaks subroutine with the default settings (−style factor -o auto). The threshold was set at a false discovery rate (FDR) of 0.001 determined by peak finding using randomized tag positions in a genome with an effective size of 2×10^9^ bp. For CTCF differential peak discovery, a combined list of peaks was generated using the HOMER command mergePeaks -d 100 with peaks called in vehicle and H2O2 condition. To find peaks that are differentially enriched between the two conditions, HOMER get Differential Peaks was used with option -F4 -P 0.0001 (for finding peaks that have 4-fold more tags and a cumulative Poisson p-value less than 0.0001, respectively). Bedgraph files were generated using HOMER scripts makeUCSCfile and makeMultiWigHub.pl for visualization in the UCSC genome browser. The total number of mappable reads was normalized to 10^7^ for each experiment presented in this study. Heatmaps were generated using deeptools2 ^87^ (version 3.5.1).

### Run-On Sequencing

Global Run On (GRO)-Seq experiments were performed as previously reported ^79^. Briefly, ∼10 million cells were washed three times with cold PBS and then sequentially swelled in swelling buffer (10 mM Tris/HCl pH7.5, 2 mM MgCl_2_, 3 mM CaCl_2_) for 5 minutes on ice, then lysed in lysis buffer (swelling buffer plus 0.5% NP-40 and 10% glycerol). The resultant nuclei were washed one more time with 10 ml lysis buffer and finally resuspended in 100 µl of freezing buffer (50 mM Tris/HCl pH 8.3, 40% glycerol, 5 mM MgCl_2_, 0.1 mM EDTA). For the run-on assay, re-suspended nuclei were mixed with an equal volume of reaction buffer (10 mM Tris/HCl pH 8.0, 5 mM MgCl_2_, 1 mM DTT, 300 mM KCl, 20 units of SUPERase-IN, 1% sarkosyl, 500 μM ATP, GTP, and Br-UTP, 2 μM CTP) and incubated for 5 min at 30°C. The resultant nuclear-run-on RNA (NRO-RNA) was then extracted with TRIzol LS reagent (Invitrogen) following the manufacturer’s instructions. NRO-RNA was fragmented to ∼300-500nt by alkaline base hydrolysis on ice and followed by treatment with DNase I and Antarctic phosphatase. These fragmented Br-UTP labeled nascent RNA was then immunoprecipitated with anti-BrdU agarose beads (sc-32323AC, Santa Cruz Biotechnology) in binding buffer (0.5x SSPE,1 mM EDTA, 0.05% Tween-20) for three hours at 4°C with rotation. Purified RNA was treated with PNK4 before being used for cDNA synthesis using NEBNext® Multiplex Small RNA Library Prep Set for Illumina® Kit (NEB). Obtained cDNA template was amplified by PCR using the LongAmp® Taq 2XMaster Mix (NEB) for sequencing.

Sequencing reads were inspected for quality control using FASTQC (https://www.bioinformatics.babraham.ac.uk/projects/fastqc/), and sequencing adaptors were trimmed, if necessary, using TRIMMOMATIC ^85^ (http://www.usadellab.org/cms/?page=trimmomatic). Reads were aligned to hg19 with Bowtie2 (version 2.26) using --very-sensitive setting. Tag directories were then generated using HOMER ^86^ (version 4 10.3), keeping only unique aligned reads per genome position but allowing up to 3 unique reads per position (−tbp 3). For read counting, the HOMER script analyzeRepeats.pl was used to estimate the raw counts per gene and also to compute the pausing ratio. The aligned reads were counted over the RefSeq gene bodies (after excluding a TSS 400bp-proximal region downstream of TSS up to 13Kb of the gene body). EdgeR (http://www.bioconductor.org/) was used to compute the significance of the differential gene expression (FC≥1.5, FDR≤0.05). Additionally, a read density threshold (i.e., GRO-seq normalized read counts/kb) was used to exclude lowly expressed genes. Bedgraph files were generated using HOMER scripts makeUCSCfile and makeMultiWigHub.pl for visualization in the UCSC genome browser. Heatmaps were generated using deeptools2 ^87^ (version 3.4.3).

### RNA-seq

RNA-seq data were generated as previously described ^88^. In brief, total RNA from left ventricle of 8-month-old mice was collected with TissueLyser II (Qiagen) and RNA was extracted using the RNeasy fibrous tissue isolation kit (Qiagen, 74704), the quality was assessed based on RNA integrity number (RIN) using an Agilent Bioanalyzer. Libraries were prepared using the Illumina TruSeq stranded mRNA kits and sequenced using Illumina NovaSeq 6000. Samples were sequenced to an average of around 25 million read pairs.

### 4C-seq

The 4C-seq experiments were conducted following a published protocol ^79^ with modification. In brief, 10 million cells were cross-linked with 1% formaldehyde for 10 min and nuclei were extracted. Nuclei were resuspended in restriction enzyme buffer and incubated with 0.3% SDS for 1 h at 37 °C and further incubated with 2% Triton X-100 for 1 h; then 400 U of DpnII restriction enzyme was added and nuclei were incubated overnight. Restriction enzyme was heat-inactivated at 65 °C for 20 min. Ligation of DNA regions in close physical proximity was performed using 1,000 U of T4 DNA ligase (NEB) for overnight. After de-cross-linking, the second digestion and ligation were performed using restriction enzyme NlaIII and T4 DNA ligase. The 4C-seq libraries were amplified using PCR with the first primer designed on each viewpoint and the second primer designed beside the NlaIII site. Both primers contained Illumina sequencing adaptors and barcode (**Suppl. Table 7**). The 4C libraries were sequenced on the Illumina HiSeq 2500 using single-read 100-cycle runs.

Analysis of 4C-seq data was done using an existing 4C pipeline ^89^. Valid reads of genomic regions were generated by clipping out the primer sequences in the raw reads. These clipped reads with unique 3′ ends were aligned to hg19 human genomic coordinates. All the samples were quality checked according to the following: ratio of the read number of cis-interactions versus the read number of total-interaction is larger than 40%. For data visualization, a 5 kb window size was chosen to compute the trend curve and the grey band on top of each 4C heat map displays the 20–80% for the windows. Median contact intensities were depicted as color-coded multi-scaled heat maps, ranging from a 2kb sliding window at the top of the heat map to a 50-kb sliding window at the bottom of the heat map.

### Overlap Analysis of Genes that gain CTCF Binding upon H_2_O_2_ Treatment

A list of all hg19 protein-coding genes was downloaded from Gencode. Differential gene expression analysis using DESeq2 was carried out to identify genes that responded to H_2_O_2_. H2O2-responsive gene promoters were defined as the regions surrounding annotated transcription start sites (−2kb/+2kb). CTCF peaks were then overlapped with gene promoters to determine those that gained CTCF occupancy upon H_2_O_2_ treatment using bedops with the “--element-of 1” parameter specified.

### Generation of *Sirt1* promoter mutant mice

For generation of Sirt1-promoter mutant transgenic mice, a microinjection of a Cas9 protein/tracrRNA/crRNA mix into mouse zygotes was performed as previously described ^90^. Briefly, an injection mix containing Cas9 protein, tracrRNA and crRNA (all: Alt-R CRISPR-Cas9, IDT, 1072532, 1081060) and ssODN at a 0.6 µM equimolar concentration in TE pH 7.5 (IDT) was prepared and incubated for 5 min at room temperature. After centrifugation at 10000 rpm for 1 min the supernatant was transferred into a new tube, stored on ice and used for injection within 1 hour. Pups were genotyped at 2 months of age. Briefly, a 0.5 cm tail clip was incubated in 50mM Tris/HCl pH 8.0; 100mM EDTA; 100mM NaCl; 1% SDS; 0.5mg/ml Proteinase K overnight at 55°C, followed by adding 100 µl 5M NaCl for 10 min and centrifugation at 15000 rpm for 15 min at 4°C for protein precipitation. The pellet was discarded and genomic DNA was recovered from the supernatant by isopropanol precipitation. A promoter region in the Sirt1 promoter was amplified by PCR (Terra PCR Direct Red Dye Premix, Takara), the PCR product gel purified (Gel Extraction Kit, Qiagen) and Sanger-sequenced (Retrogen). The abi tracks were inspected for SNPs at the expected positions and called as wild-type, hetero- and homozygous manually.

### Echocardiography

Echocardiography (ECG; M-mode, two-dimensional, tissue Doppler and, pulse-wave Doppler) was performed under isoflurane anesthesia (1%–1.5% at a flow rate of 1 l/min oxygen) using a small-animal, high-resolution Vevo 3100 imaging unit (MX550S 26–52 MHz transducer, FUJIFILM VisualSonics, Toronto, Canada). For the assessment of systolic function and tissue Doppler analysis, ECG-traced heart rates were maintained between 520 and 600 beats/min to avoid anesthesia-induced changes in cardiac function. To assess systolic cardiac function, left ventricular dimensions, fractional shortening (%FS), mean circumferential fiber shortening rate (VCF), and wall thickness were measured, as previously described ^91,92^.

### Cardiac myocyte isolation

Neonatal mouse ventricular myocytes (NMVMs) were isolated from 0-to 2-day-old mouse hearts as previously described ^88^. Briefly, hearts were excised, the atria were removed and then the ventricular pieces were digested overnight in 50 ml of 0.5 mg/ml trypsin-HBSS at 4°C. On the following day, 25 ml of the trypsin solution was replaced by serum-free medium and digested for 3-4 minutes with gentle agitation (100-150 rpm). Subsequently the trypsin/serum free medium solution was removed, and digestion was continued in type II collagenase (200-300 Units/ml, Worthington Biochemical Corporation) for three additional digestions, each for 3-5 min. Adding ice-cold serum-containing medium to inactivate collagenase-II terminated each collagenase digestion. Supernatants from digests were pooled and cells were collected by centrifugation. Two pre-plating steps, each for 1 h on uncoated tissue-culture dishes, were used to further purify NMVMs from this supernatant. Only non-myocyte cells rapidly adhere to these uncoated dishes. Approximately 5×10^6^-6×10^6^ cells were plated onto each 60-mm culture dish, which were coated with 0.2% gelatin/fibronectin (1 µg/ml; F0635, Sigma) to generate confluent beating cultures. Cells were maintained in serum-containing medium (75% DMEM, 25% M-199, 10% fetal bovine serum and 5% horse serum). One week after plating, only confluent and beating cultures of NMVMs were used.

### Single nucleus RNA-seq of murine heart

Nuclei isolation from adult mouse hearts was performed as previously described with minor modifications ^93^. Briefly, after proper anesthesia and thoracotomy, mouse heart was harvested and placed in cold 1x phosphate buffered saline (PBS) to remove excess blood. The tissue was then minced thoroughly into tiny pieces in a clean empty petri dish over ice and 2 ml of ice-cold lysis buffer was added (0.32 M sucrose, 5 mM CaCl_2_, 3mM magnesium acetate, 2.0 mM EDTA, 0.5 mM EGTA, 10 mM Tris-HCl (pH 8.0), 1 mM DDT, 0.2% Triton X-100) supplemented with protease inhibitor cocktail (Sigma). Next, the mixture was dounced in a glass douncer in ice (7 ml, Kimble D9063, Sigma) with 50 strokes using the type A pestle followed by 3 strokes with type B pestle. The solution was filtered through a 70 and 10 μm nylon cell strainers (Falcon) attached to a 50 ml tube. The tubes were centrifuged for 5 minutes at 1000g at 4° C, and the supernatant was carefully removed and discarded. Nuclei pellet was washed three times in 5 ml of resuspension buffer (1x phosphate buffered saline, 1% BSA) supplemented with 0.05 U/µl RNase inhibitor (AM2696, Invitrogen). Last wash was performed into a 5 ml tube and spin for 5 minutes at 1000g at 4° C. Nuclei were resuspended in 300 µl of resuspension buffer supplemented with 0.2 U/µl RNase inhibitor. Integrity and Precise yield of nuclei was assessed using DAPI staining, followed by counting in chamber slide using Countess II Automated Cell Counter (Invitrogen). Resuspension buffer was added to the nuclei suspension to achieve a final concentration of 700-1200 nuclei/µl. A total of 15k nuclei were used for droplet formation in the 10X Genomics Chromium controller according to the manufacturer’s instructions in the Chromium Single Cell 3′ Reagent Kit v.3.1 User Guide. Additional components used for library preparation include the Chromium Next GEM Single Cell 3′ Kit v.3.1 (PN-1000269), the Chromium Next GEM Chip G (PN-1000127) and Dual Index Kit TT Set A (PN-1000215). Final libraries were sequenced on NovaSeq (Illumina, software v1.5) to a mean read depth of at least 50,000 total aligned reads per nuclei.

### Analysis of the single nuclei RNA-seq

Raw sequencing reads were aligned to the mouse mm10 genome using the Cell Ranger v.6.0.1 pipeline from 10X Genomics which generate a gene-barcode matrix. Seurat was used for the downstream analysis. Low-quality nuclei having high mitochondria contamination, low number of UMI and limited number of genes were discarded. RNA ambient contamination was corrected using soupX and doublets were identified and removed using DoubletFinder. After the QC steps, we normalized the data (NormalizeData function, 10,000 default scale factor) and performed a linear regression on all genes (ScaleData function). We then performed a linear dimensional reduction (RunPCA function) and batch correction using Harmony. Significant principal components were used for downstream clustering into distinct populations using the FindClusters function and uniform manifold approximation and projection (UMAP) dimensionality reduction was used for visualization purpose. We defined cardiac cell-types using canonical marker genes predominantly expressed in the different clusters. We then isolated specific clusters (WhichCells function) for subsequent analysis on the cardiomyocytes and fibroblast populations. DEGs were computed for each cell-type to test significantly differentially expressed genes between wild-type and Sirt1-Mut using the function FindMarkers. For visualization of gene expression data between different samples we used different Seurat functions including FeaturePlot, VlnPlot and DotPlot.

### Data availability

Next-gen sequencing data sets generated from this study can be accessed at GEO using accession ID GSMXXXXXXX-GSMXXXXXXX.

---

**Supplementary Figure 1.**
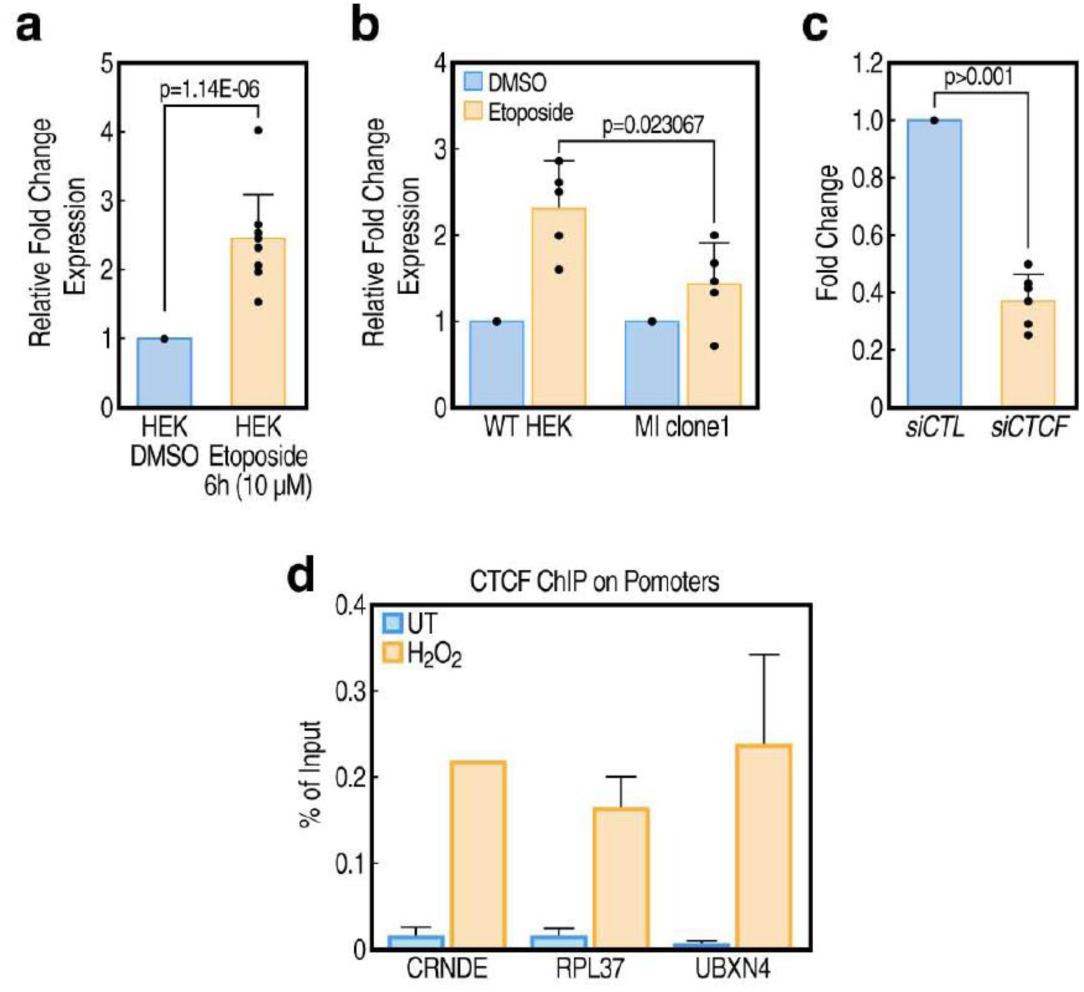
(a) SIRT1 mRNA levels measured by qPCR in HEK293 cells after 6h of 10 μM etoposide. Fold change expression of SIRT1 normalized to GAPDH and compared to untreated cells (UT) as shown; n=8; p-value = Student’s t-test. (b) CRISPR/Cas9 engineered HEK293 cells carrying the −92A>G mutation rs3740053 (MI clone 1) treated with 10 μM etoposide as described in pane (a), expression was measured by qPCR compared to isogenic control HEK293 cells (WT HEK) (n=4, +/-s.d.; p-value Student’s t-test). (c) Cells transfected with siRNA targeting CTCF show reduced CTCF mRNA expression compared to non-targeting control siRNA. (d) CTCF ChIP-qPCR on the *CRNDE*, *RPL37*, and *UBXN4* genes promoter in UT or after H_2_O_2_ treatment in HEK293 cells.

**Supplementary Figure 2.**
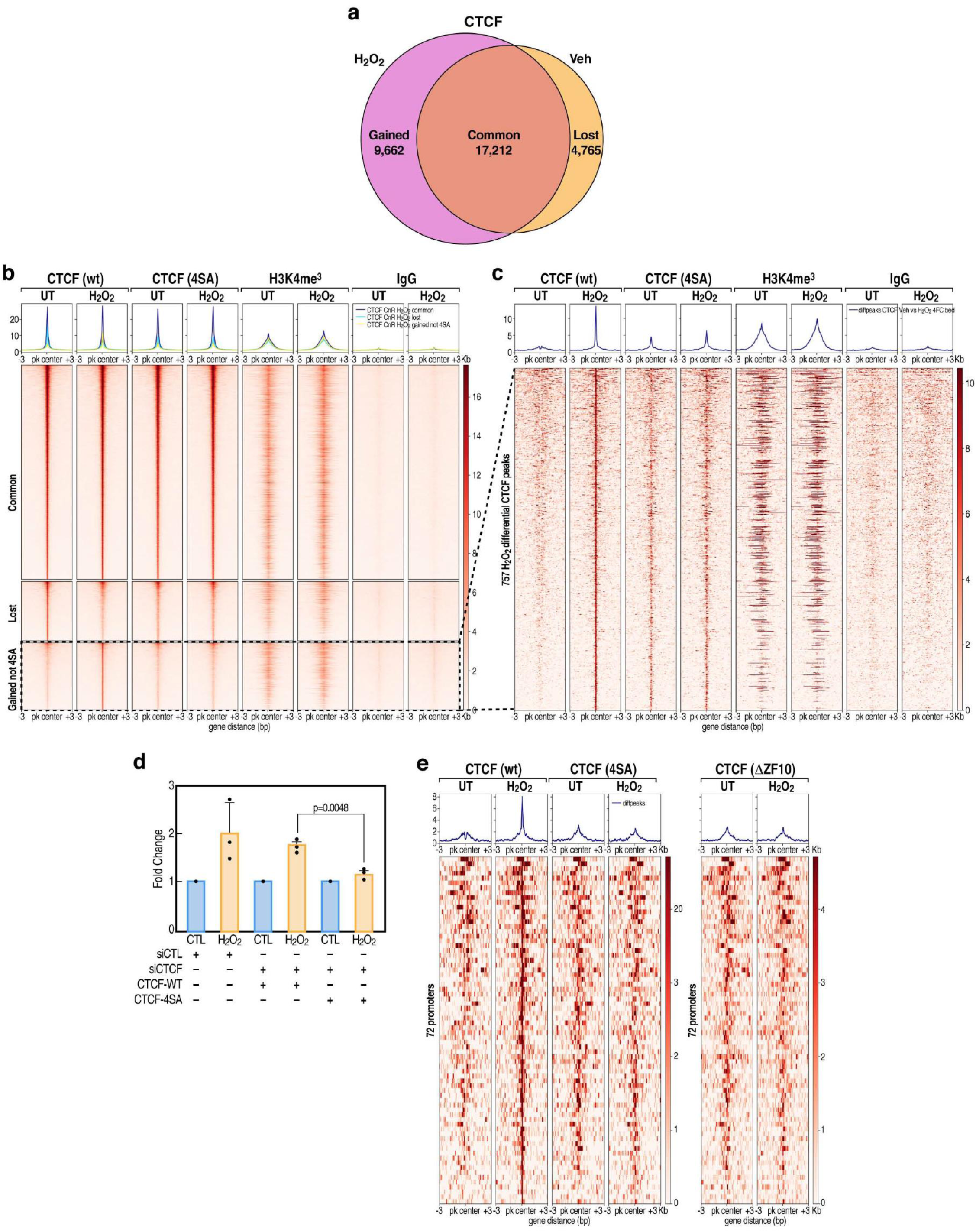
(a) Venn diagram represents the total number of CTCF peaks identified by CUT&RUN in UT or H_2_O_2_ treated HKE293 cell line. (b) Heatmap representing different numbers of CTCF peaks identified in common, lost, or gain in 4SA (CTCF mut) or CTCF-WT upon H_2_O_2_ treatment in Flag CUT&RUN. (c) Expanded view of Flag-CTCF peaks which were lost in 4SA CTCF mutant upon H2O2 induction. (d) Endogenous CTCF of HEK293 cells was depleted by 3’-UTR targeting siRNA and reconstituted with either Flag-tagged CTCF-WT or CK2-phosphorylation incompetent CTCF-4SA (S606,609,610,612A). After stimulation with 0.5 mM H_2_O_2_ SIRT1 expression was quantified by qPCR. (e) Endogenous CTCF of HEK293 cells was depleted by 3’-UTR targeting shRNA and reconstituted with Flag-tagged CTCF-WT or CTCF-4SA or CTCF-ZnF10 deletion. Heatmap shows 72 promoters that show increased binding of CTCF upon H2O2 treatment (0.5 mM H_2_O_2_ 15 min, 2 hours of wash-off) while lost to bind by CTCF-4SA or CTCF-ZnF10 deletion.

**Supplementary Figure 3.**
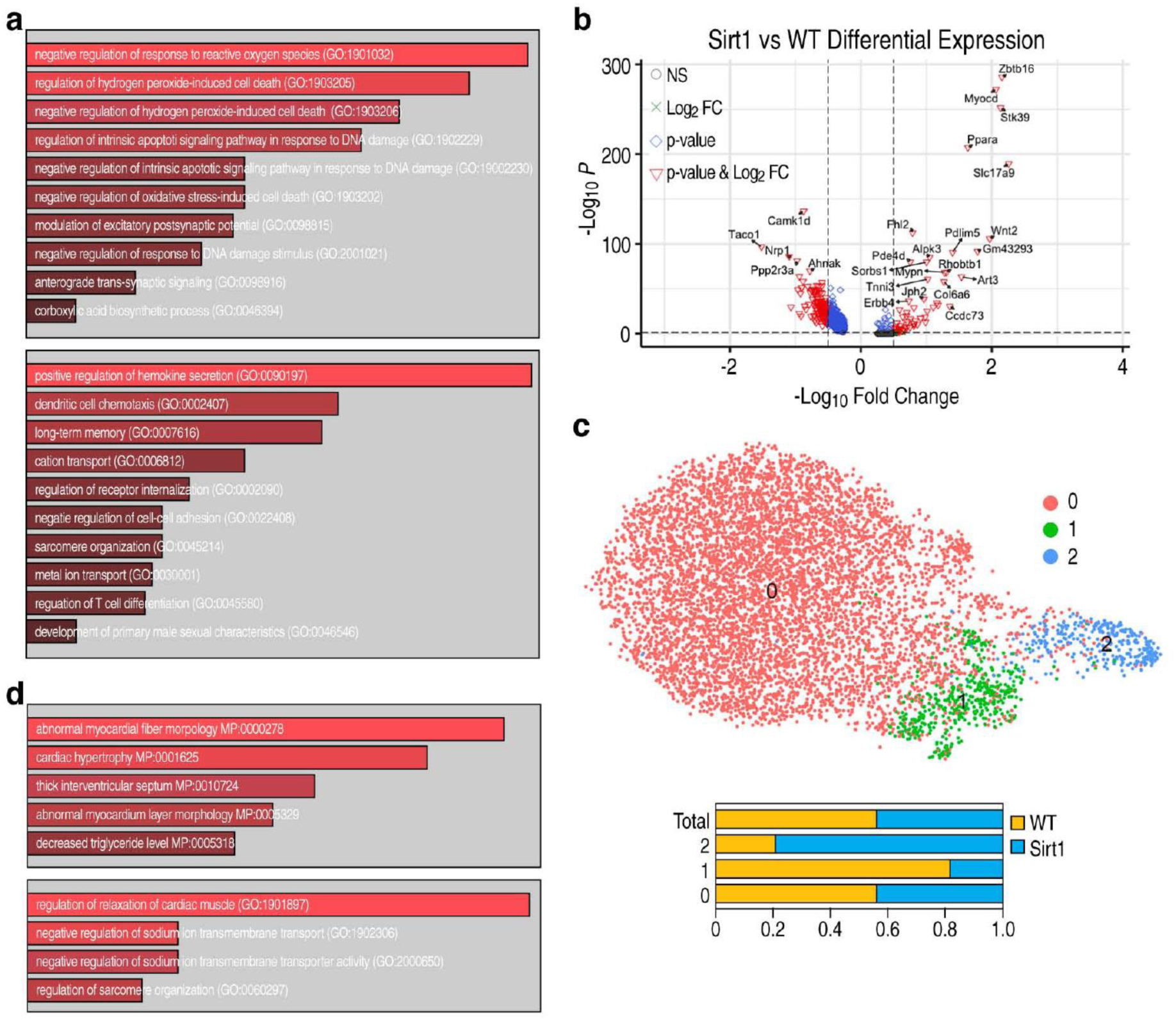
(a) Gene Ontology (GO) terms for differentially expressed genes identified in bulk RNA-seq were performed on 10 WT promoter male mice and 12 Sirt1-promoter mutant male mice. (b) Volcano plot of differentially expressed genes across all cell types identified in snRNA-seq performed in duplicates of wild-type control and Sirt1-Mut mice. (c) UMAP analysis representing that almost 50 percentage of differentially expressed genes belong to cluster 0, cardiomyocytes. (d) GO terms for the differentially expressed genes (down-, top panel; and up-regulated, bottom panel, respectively) identified in snRNA-seq as described in panel (b).

**Supplementary Figure 4.**
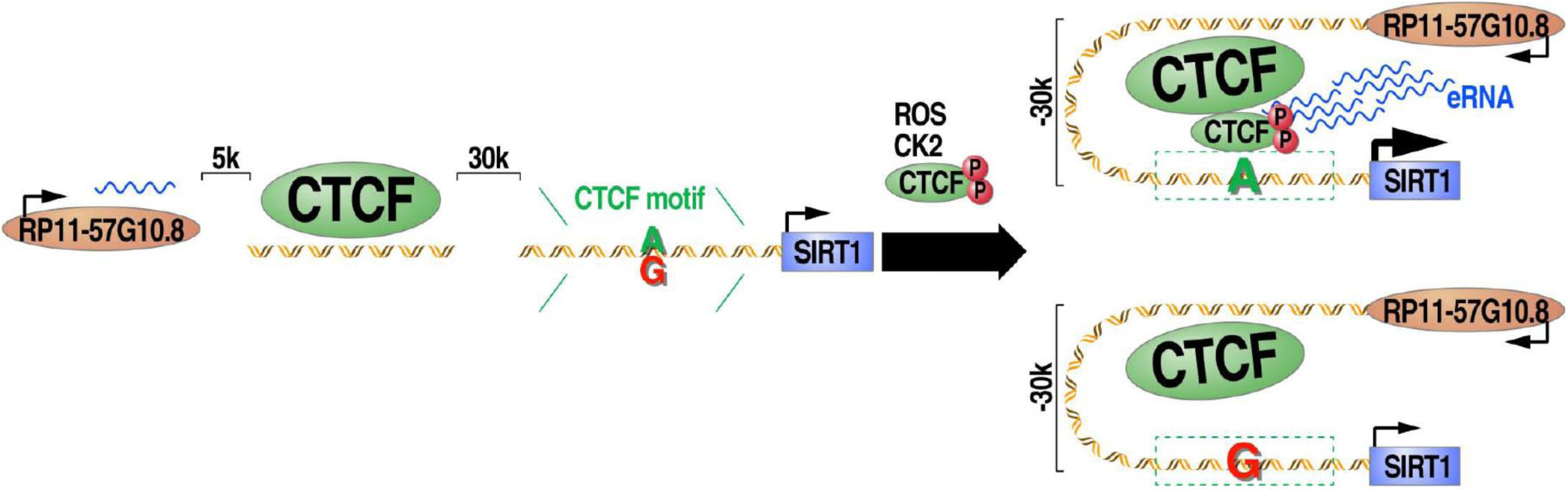
Model of CTCF regulation of SIRT1 gene transcription in response to oxidative stress. CTCF is phosphorylated by CK2 triggered by ROS, with CTCF binding to a low affinity binding site in the SIRT1 promoter. CTCF binding leads to a local change in 3D genome architecture, leading to interaction of an upstream lncRNA region to the SIRT1 promoter, with the lncRNA plausibly acting as an enhancer for SIRT1. This regulation is lost when the CTCF binding sites in the SIRT1 promoter is further affected by a single nucleotide variant A→G at the −92 position of the SIRT transcriptional start site, inhibiting CTCF binding to cardiomyocytes SIRT1 promoter, and resulting in defective cardiac systolic relaxation *in vivo*.

