## Supplemental Table 1-4 for "Recruitment of CTCF to the SIRT1 promoter after Oxidative Stress mediates Cardioprotective Transcription"

**Supplement Table 1.**
**Association of *SIRT1* tagSNPs with MI in SALSA (N=356).**

| SNP (A/a)[a] | Position[b] | Location | MAF[c] Case | MAF[c] Control | *P* |
|---|---|---|---|---|---|
| rs3740051 (A/G) | 69313965 | IVS 1-468 | 0.36 | 0.12 | **0.016** |
| rs2236318 (T/A) | 69318575 | IVS 3-71 | 0.18 | 0.12 | **0.004** |
| rs2236319 (A/G) | 69319226 | IVS 3+339 | 0.36 | 0.13 | **0.016** |
| rs10823103 (G/A) | 69320975 | IVS 4-191 | 0.29 | 0.13 | **0.003** |
| rs7896005 (A/G) | 69321131 | IVS 4-35 | 0.21 | 0.13 | **0.004** |
| rs1885472 (C/G) | 69324818 | IVS 4+3500 | 0.29 | 0.13 | **0.004** |
| rs10997866 (A/G) | 69328657 | IVS 4+7339 | 0.25 | 0.13 | **0.008** |
| rs10823107 (A/G) | 69004827 | IVS 5-6323 | 0.42 | 0.13 | 0.059 |
| rs10823108 (G/A) | 69330500 | IVS 5-6053 | 0.36 | 0.13 | **0.014** |
| rs7069102 (G/C) | 69333126 | IVS 5-3427 | 0.29 | 0.13 | **0.004** |
| rs7096385 (C/T) | 69334887 | IVS 5-1666 | 0.36 | 0.13 | **0.018** |
| rs2224573 (A/G) | 69335393 | IVS 5-1160 | 0.31 | 0.13 | **0.018** |
| rs2273773 (T/C) | 69336604 | Ex5+97 (L332L) | 0.36 | 0.13 | **0.015** |
| rs10823109 (T/G) | 69336951 | IVS 5+251 | 0.38 | 0.13 | **0.011** |
| rs10997870 (G/T) | 69338020 | IVS 6+132 | 0.21 | 0.13 | **0.003** |
| rs10823111 (T/C) | 69340525 | IVS 7+1320 | 0.31 | 0.13 | **0.011** |
| rs10823112 (A/G) | 69340822 | IVS 8-1415 | 0.36 | 0.12 | **0.015** |
| rs1467568 (A/G) | 69345164 | IVS 9-864 | 0.21 | 0.13 | **0.004** |
| rs4746720 (T/C) | 69346836 | 3'UTR +809 | 0.07 | 0.13 | 0.870 |
| rs10997875 (C/T) | 69349830 | IVS 9+1683 | 0.29 | 0.13 | **0.004** |

(A/a)[a], major allele/minor allele; Position[b], NCBI human genome build 36 coordinates; MAF[c], minor allele frequency; OR[d], based on minor(or major) allele (95% CI)

**Supplement Table 2.**
**Replication of *SIRT1* gene association with MI in SAFHS (N=980)**

| SNP (A/a) | Position | *P* |
|---|---|---|
| rs17712705 (A>G) | 69293277 | $1.56 \times 10^{-4}$ |
| rs12778366 (T>C) | 69313085 | 0.578 |
| rs2236319 (A>G) | 69319226 | 0.215 |
| rs10823108 (G>A) | 69330500 | $1.75 \times 10^{-4}$ |
| rs2273773 (T>C) | 69336604 | 0.216 |

**Supplement Table 3.**
**Sequencing analysis of *SIRT1* gene region in MI and controls.**

| Nature of variation | | rs number | Frequency | |
|---|---|---|---|---|
| location | sequence change | | with MI (N=14) | without MI (N=15) |
| Ex1-4004 | A>C | rs4746716 | 0.25 | 0.50 |
| Ex1-4000 | C>T | rs117977601 | 0.21 | 0.50 |
| Ex1-3696 | del A | rs80249390 | 0.07 | 0.13 |
| Ex1-3310 | del C | rs145136053 | 0.32 | 0.58 |
| Ex1-3058 | C>A | rs10997854 | 0.29 | 0.58 |
| Ex1-3026 | G>A | rs10997855 | 0.29 | 0.58 |
| Ex1-2953 | C>T | NA | 0.00 | 0.03 |
| Ex1-2888 | C>T | rs142194353 | 0.18 | 0.08 |
| Ex1-2884 | G>C | NA | 0.00 | 0.03 |
| Ex1-2777 | del TT | NA | 0.04 | 0.00 |
| Ex1-2734 | G>C | rs10823100 | 0.25 | 0.15 |
| Ex1-2721 | T>C | rs10997856 | 0.29 | 0.54 |
| Ex1-2716 | T>A | rs10823101 | 0.43 | 0.12 |
| Ex1-2712 | T>A | rs10997857 | 0.43 | 0.12 |
| Ex1-2594 | T>A | rs10823102 | 0.29 | 0.53 |
| Ex1-2473 | C>T | rs12250285 | 0.25 | 0.57 |
| Ex1-2369 | ins T | rs148743102 | 0.07 | 0.03 |
| Ex1-1759 | T>A | rs10740280 | 1.00 | 1.00 |
| Ex1-1348 | T>C | rs12778366 | 0.21 | 0.13 |
| Ex1-1085 | C>T | rs3758391 | 0.29 | 0.53 |
| Ex1-956 | G>T | NA | 0.04 | 0.00 |
| Ex1-810 | A>C | rs35706870 | 0.07 | 0.03 |
| Ex1-468 | A>G | rs3740051 | 0.39 | 0.23 |
| Ex1-210 | A>C | rs932658 | 0.29 | 0.53 |
| Ex1-92 | A>G | rs3740053 | 0.36 | 0.20 |
| Ex1-86 | G>C | rs2394443 | 0.21 | 0.53 |
| 5'UTR 35(-18) | C>T | NA | 0.04 | 0.00 |
| 5'UTR 39(-14) | A>G | NA | 0.04 | 0.00 |
| IVS2+41 | G>T | rs932657 | 1.00 | 1.00 |
| IVS2+105 | C>T | NA | 0.07 | 0.00 |
| 994(332L>L) | T>C | rs2273773 | 0.36 | 0.20 |
| IVS6+132 | G>T | rs10997870 | 0.46 | 0.27 |
| 3UTR+209 | insT | rs61666042 | 0.43 | 0.36 |
| 3UTR+209 | A>T | rs2394445 | 0.29 | 0.20 |
| 3UTR+263 | del TTTC | rs35461348 | 0.21 | 0.30 |
| 3UTR+479 | C>T | rs4746720 | 0.11 | 0.07 |
| 3UTR+710 | C>T | rs752578 | 1.00 | 1.00 |
| 3UTR+1109 | del | NA | 0.04 | 0.00 |

**Supplement Table 4.**
**JASPAR motif analysis of a 100 nt fragment surrounding SNP rs3740053 at the -92 position of the *SIRT1* promoter.**

| # | Matrix ID | Name | Score | Relative score | Sequence ID | Start | End | Strand | Predicted sequence# |
|---|---|---|---|---|---|---|---|---|---|
| 1A# | MA0139.1 | MA0139.1. CTCF | 2.4438772 | 0.7312807897765095 | hg19_dna | 46 | 64 | + | G**A**GCGCCGAG AGGGCGGGG |
| 1G# | MA0139.1 | MA0139.1. CTCF | 3.7159605 | 0.7460492553770443 | test minority G | 46 | 64 | + | G**G**GCGCCGAG AGGGCGGGG |
| 2 | MA0139.1 | MA0139.1. CTCF | 1.4640013 | 0.7199047548507604 | hg19_dna | 4 | 22 | - | ACAACACTAC GGGTCACGT |
| 3 | MA0139.1 | MA0139.1. CTCF | -0.46506122 | 0.6975089779437107 | hg19_dna | 52 | 70 | + | CGAGAGGGCG GGGGCGGCG |
| 4 | MA0139.1 | MA0139.1. CTCF | -2.183274 | 0.677561096248453 | hg19_dna | 63 | 81 | + | GGGCGGCGAT GGGGCGGGT |
| 5 | MA0139.1 | MA0139.1. CTCF | -2.431107 | 0.6746838370426304 | hg19_dna | 78 | 96 | + | GGGTCACGTG ATGGGGTT |
| 6A# | MA0139.1 | MA0139.1. CTCF | -2.4947338 | 0.6739451513599255 | hg19_dna | 29 | 47 | + | GCCCGCGTGG GTGGCGGG**A** |
| 6G# | MA0139.1 | MA0139.1. CTCF | -3.076976 | 0.6671855116686177 | test minority G | 29 | 47 | + | GCCCGCGTGG GTGGCGGG**G** |
| 7 | MA0139.1 | MA0139.1. CTCF | -3.2310648 | 0.6653965924821084 | hg19_dna | 27 | 45 | + | TGGCCCGCGT GGGTGGCGG |
| 8 | MA0139.1 | MA0139.1. CTCF | -4.1206765 | 0.6550684950367299 | hg19_dna | 49 | 67 | + | CGCCGAGAGG GCGGGGGCG |

JASPAR (https://jaspar.genereg.net) motif analysis of a 100 nt fragment of the CTCF promoter (hg19_dna range=chr10:69644289-69644388) including SNP rs3740053 with at the -92-position relative to the SIRT1 transcriptional start site with CTCF PWM (MA0139.1) using a relative profile score threshold of 65%.
### SNP rs3740053 highlighted in bold
