## Supplemental Table 6 for "Recruitment of CTCF to the SIRT1 promoter after Oxidative Stress mediates Cardioprotective Transcription"

**Supplement Table 6.**
**Echocardiography of wild-type (wt) and *Sirt1*-promoter mutant mice (Sirt1-Mut).**

| ♀ | 4-month-old | | | 5-month-old | | | 7-month-old | | |
|---|---|---|---|---|---|---|---|---|---|
| **Parameters** | **wt** | **Sirt1-Mut** | **p-value** | **wt** | **Sirt1-Mut** | **p-value** | **wt** | **Sirt1-Mut** | **p-value** |
| **HR** (bpm) | 569 ± 41 | 572 ± 39 | ns | 615 ± 56 | 599 ± 32 | ns | 600 ± 57 | 586 ± 34 | ns |
| **IVSd** (mm) | 0.67 ± 0.03 | 0.68 ± 0.06 | ns | 0.67 ± 0.04 | 0.63 ± 0.05 | ns | 0.66 ± 0.08 | 0.67 ± 0.04 | ns |
| **LVPWd** (mm) | 0.67 ± 0.04 | 0.68 ± 0.06 | ns | 0.68 ± 0.01 | 0.65 ± 0.04 | ns | 0.68 ± 0.03 | 0.70 ± 0.03 | ns |
| **LVMass** (mg) | 75.4 ± 9.1 | 73.6 ± 13.3 | ns | 68.2 ± 8.8 | 73.4 ± 7.9 | ns | 69.6 ± 7.3 | 75.3 ± 23.6 | ns |
| **LVIDd** (mm) | 3.53 ± 0.23 | 3.41 ± 0.34 | ns | 3.28 ± 0.22 | 3.58 ± 0.27 | p=0.03 | 3.38 ± 0.23 | 3.64 ± 0.25 | p=0.04 |
| **LVIDs** (mm) | 2.36 ± 0.20 | 2.59 ± 0.37 | ns | 2.15 ± 0.16 | 2.77 ± 0.30 | p=0.0001 | 2.27 ± 0.29 | 2.90 ± 0.31 | p=0.0004 |
| **%FS** | 33.2 ± 2.80 | 24.1 ± 5.64 | p=0.001 | 34.4 ± 2.41 | 22.7 ± 4.16 | p<0.0001 | 33.1 ± 4.24 | 20.3 ± 3.95 | p<0.0001 |
| **BW** (g) | 22.0 ± 1.0 | 21.9 ± 1.2 | ns | 23.0 ± 1.3 | 23.1 ± 1.2 | ns | 23.5 ± 1.3 | 23.3 ± 1.1 | ns |
| **n =** | 7 | 12 | | | | | | | |

| ♂ | 4-month-old | | | 5-month-old | | | 7-month-old | | |
|---|---|---|---|---|---|---|---|---|---|
| **Parameters** | **wt** | **Sirt1-Mut** | **p-value** | **wt** | **Sirt1-Mut** | **p-value** | **wt** | **Sirt1-Mut** | **p-value** |
| **HR** (bpm) | 558 ± 61 | 560 ± 46 | ns | 593 ± 51 | 568 ± 56 | ns | 569 ± 43 | 585 ± 30 | ns |
| **IVSd** (mm) | 0.74 ± 0.04 | 0.71 ± 0.04 | ns | 0.70 ± 0.06 | 0.66 ± 0.02 | p<0.0001 | 0.75 ± 0.05 | 0.69 ± 0.04 | p=0.005 |
| **LVPWd** (mm) | 0.71 ± 0.02 | 0.69 ± 0.03 | ns | 0.71 ± 0.03 | 0.66 ± 0.04 | p=0.01 | 0.73 ± 0.04 | 0.67 ± 0.01 | p=0.0002 |
| **LVMass** (mg) | 83.3 ± 14.2 | 84.1 ± 9.5 | ns | 73.1 ± 13.7 | 82.2 ± 9.9 | ns | 89.6 ± 29.2 | 90.4 ± 10.7 | ns |
| **LVIDd** (mm) | 3.52 ± 0.38 | 3.63 ± 0.24 | ns | 3.33 ± 0.35 | 3.71 ± 0.36 | p=0.02 | 3.76 ± 0.34 | 3.87 ± 0.26 | ns |
| **LVIDs** (mm) | 2.34 ± 0.30 | 2.86 ± 0.26 | p=0.0003 | 2.20 ± 0.29 | 2.94 ± 0.29 | p<0.0001 | 2.59 ± 0.24 | 3.06 ± 0.23 | p<0.0001 |
| **%FS** | 33.50 ± 2.73 | 21.22 ± 0.26 | p<0.0001 | 33.93 ± 4.01 | 20.65 ± 3.49 | p<0.0001 | 31.05 ± 4.76 | 20.92 ± 2.63 | p<0.0001 |
| **BW** (g) | 26.1 ± 1.7 | 26.3 ± 2.2 | ns | 25.7 ± 1.9 | 26.1 ± 2.3 | ns | 27.4 ± 1.9 | 27.8 ± 2.2 | ns |
| **n =** | 10 | 12 | | | | | | | |

HR – heart rate; IVSd – interventricular septal end diastole; LVPWd – left ventricular posterior wall end diastole; LVMass – left ventricle mass; LVIDd/s – left ventricular internal diameter end diastole/end systole; %FS – percent fractional shortening; BW – whole body weight
p-values: Student’s t-test, ns = not significant (p>0.05)
